# Tumor Landscape Analysis: An Ecologically Informed Framework to Understand Tumor Microenvironments

**DOI:** 10.1101/2025.04.22.646608

**Authors:** Luis Cisneros, Merih Deniz Toruner, Martin E. Fernandez-Zapico, Carlo C. Maley, Ryan M. Carr

## Abstract

Tumor microenvironments (TMEs) are spatially complex and dynamic systems shaped by evolutionary pressures, tissue architecture, and cellular interactions. To capture this complexity, we developed the Tumor Landscape Analysis (TLA) pipeline, a computational framework applying principles from landscape ecology and spatial statistics to quantitatively characterize tumor spatial heterogeneity. TLA integrates spatially resolved pathology data, including whole-cell, point-based, and region-level formats, and computes multiscale metrics to assess cell distributions, neighborhood relationships, and tissue-level organization. The framework leverages ecological indices such as the Morisita-Horn index, Ripley’s H function, and Shannon diversity to quantify intercellular proximity, spatial clustering, and cellular diversity. It also uses the concept of local microenvironments (LMEs), data-driven ecological niches defined by local cell-type abundance and spatial uniformity, enabling unsupervised and reproducible classification of tumor regions. The use of fragmentation metrics, including patch density, shape complexity, and interspersion, provide further insight into spatial disorganization and emergent tissue architecture. TLA is agnostic to imaging modality and biological context, supporting broad applicability across tumor types and sample formats. By translating complex tissue architectures into interpretable spatial metrics, the pipeline enables integrative analyses that link spatial ecology to clinical and molecular phenotypes. This approach facilitates a deeper understanding of how spatial features contribute to tumor progression, therapeutic resistance, and clinical outcomes, offering new opportunities for precision oncology rooted in spatial systems biology.

## Introduction

Cancer is a dynamic disease shaped by genetic and epigenomic variations, as well as environmental pressures driving adaptive changes within the tissue ecosystem. Tumor initiation and progression involve continuously evolving interactions among tumor cells, stromal elements, and non-cellular components of the microenvironment. Landscape ecology offers a compelling framework to interpret this spatial complexity (1). Systemic and local regulators, combined with structural constraints, promote clonal divergence, cell migration, metastasis, and the emergence of therapy-resistant subclones (2,3). Therapeutic interventions act as selective pressures, enhancing clonal diversity and resistance (4,5). Moreover, the coevolution between cancer cells and their microenvironment creates a feedback loop that sustains progression despite treatment (6,7). Viewing cancer through an eco-evolutionary lens not only deepens our understanding of tumor behavior but also opens pathways for risk stratification and treatment strategies leveraging cancer’s ecological dynamics.

The functional integrity of tissues depends on the spatial coordination of diverse cell types. Disease disrupts these interactions, compromising function and altering spatial organization—much like habitat fragmentation in ecology, which leads to biodiversity loss and systemic environmental shifts (8). This parallel underscores the value of a landscape ecology approach in cancer biology (9). The rise of spatial biology has driven demand for tools capable of analyzing cellular spatial structures and interactions. Advances in multiplexed imaging now allow high-resolution mapping of tissue complexity, enabling deep insights into the tumor microenvironment (TME) (10–12). In this context, the TME can be viewed as a multicellular ecosystem, where intercellular interactions and spatial heterogeneity critically influence tumor progression and therapeutic resistance (13).

Traditional spatial assessments rely on qualitative histopathologic review—insightful but labor-intensive and variable (14). Landscape ecology, by contrast, provides quantitative tools to analyze spatial heterogeneity through concepts such as species mixing, parasitism, predator-prey dynamics, and spatial entropy. These metrics can objectively assess cellular diversity, segregation, and interaction patterns (15), revealing non-random spatial relationships. Ecological models adapted for oncology help elucidate immune-tumor interactions, necrosis, and stromal architecture (16,17).

Increasingly, ecological metrics are used to model tumor heterogeneity and population dynamics. The Morisita-Horn index, for instance, quantifies immune-tumor colocalization with prognostic value in breast cancer (18). Computational approaches have further advanced this paradigm: Pacheco et al. (2014) employed evolutionary game theory to model cell dynamics in monoclonal gammopathy (19), while Andor et al. (2016) used tools like EXPANDS and PyClone to map clonal evolution and its correlation with clinical outcomes (20). These tools help reconstruct tumor evolutionary trajectories, offering insights into progression, resistance, and prognosis. Integrating ecological and computational metrics into oncology thus holds promise for translating spatial complexity into clinical utility.

Emerging imaging platforms enable unprecedented resolution of the tumor landscape. Studies by Heindl et al. and Keren et al. demonstrate how digital pathology and multiplex imaging reveal single-cell interactions within the TME (21–23). Three-dimensional reconstructions now illuminate invasion patterns and regional heterogeneity in greater depth (24), supporting not only visualization but also spatially resolved analysis of tumor architecture.

Despite these advances, applying ecological frameworks to cancer remains methodologically complex. Tumor tissues are inherently heterogeneous and dynamic, requiring tools capturing spatial and temporal variation. To meet this need, we developed a Tumor Landscape Analysis (TLA) pipeline, a computational framework inspired by landscape ecology and environmental systems modeling. TLA integrates spatial statistics and ecological metrics to quantify heterogeneity across scales, capturing cellular distributions, interclass relationships, and microarchitectural features within the TME. The pipeline includes data preprocessing, computation of landscape metrics, and statistical aggregation of spatial features. These outputs can be integrated with disease variables to define associations between ecological patterns and clinical phenotypes. By translating tissue architecture into interpretable spatial data, TLA provides a powerful strategy to decode tumor ecology, offering novel insights into prognosis, disease progression, and therapeutic response.

## Materials and Methods

### TLA Pipeline Architecture

The TLA pipeline is a computational framework designed to quantify spatial heterogeneity in tumor tissues by integrating concepts from landscape ecology and spatial statistics (**Figure 1**). It accommodates diverse data types, including whole-cell segmentations, cell centroid coordinates, and region-level annotations. The pipeline analyzes individual cell features, computes neighborhood-level spatial profiles, and evaluates tissue architecture using fragmentation metrics. The framework identifies local microenvironments (LMEs), which represent discrete ecological niches based on local cell-type abundance and spatial uniformity. These outputs are synthesized into interpretable spatial maps and multiscale metrics, which can be integrated with clinical and molecular data to explore associations between spatial architecture and disease phenotypes. TLA is designed to be broadly applicable across imaging platforms and cancer types, supporting a systems-level approach to tumor ecology.

**Figure 1.**
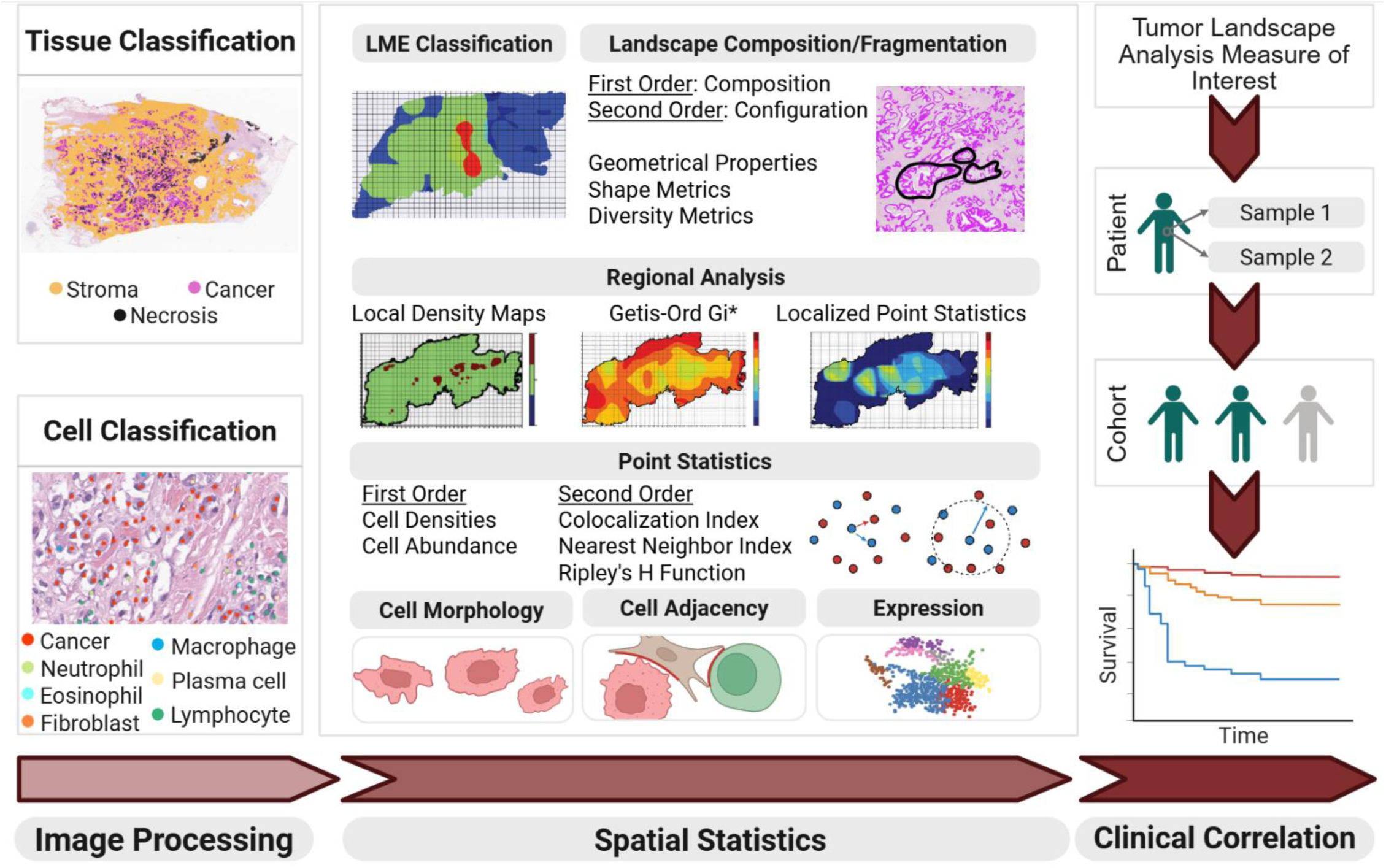
Tumor Landscape Analysis (TLA) Workflow showing the different stages of the analysis pipeline. Left panel: Tissue and Cell classification are performed using computer vision assisted by machine learning, and provides the data analyzed in TLA. Middle panel: The lower level of analysis consists of individual cell characterization. Then point statistics is done for cell coordinate data. Regional analysis includes the characterization of regions and local spatial profiles. Finally, with data provided from tissue classification, or local microenvironment (LME) classification based on local profiles, discrete patches, or niches, are defined and used for estimating fragmentation statistics. Right panel: TLA measures that are estimated at the whole-slide scale (including global spatial statistics and fragmentation statistics) are collected and summarized for patient-level analysis that includes statistical modeling, and survival analysis in combination with clinical variables and outcomes.

### Data Types and Segmentation Approaches

Accurate segmentation and classification of tissue features are foundational for spatial analysis (23,24,29,32,33). Digital pathology uses deep learning methods and multiplex imaging to generate spatially resolved maps of tissues. These computational methods generally fall into three categories based on the scale and type of segmentation performed, including the identification and classification of whole-cell images, cell nuclei locations, and multiple-cell tissue component regions.

### Cell-Level Spatial Metrics

Cell-level descriptions enable the generation of summary statistics that characterize cellular properties based on groupings defined by cell type or spatial criteria, such as membership within specific regions of interest, tissue compartments, or spatially resolved transcriptomic domains (37). These statistics are essential for quantifying localized phenotypic features relevant to tissue ecology and tumor biology. By linking individual cellular characteristics to their spatial context, such analyses facilitate detailed inferences about spatial dynamics and the relationships between cellular behaviors and tissue-level organization.

When digital pathology provides whole-cell detection data, several types of measurements can be derived directly from the properties of individual cells. Morphological features, including area, perimeter, centroid, equivalent radius, and circularity, provide information about cell shape and size. Expression characteristics can be quantified through multiplex imaging, which yields marker-specific intensity values for each cell (22,23,32). Additionally, cellular context can be assessed through adjacency analyses that quantify the number and types of neighboring cells, as well as the edge lengths shared between them.

These cell-level features form the basis for multi-scale modeling of the tumor microenvironment and spatially informed biological hypotheses. Detailed mathematical derivations for calculating equivalent radius, circularity, and adjacency edge features are provided in the **Supplementary Methods**.

### Global Point-Based Spatial Statistics

Point statistics are employed to characterize spatial features when digital pathology data are provided in the form of cell coordinates. This analytical approach focuses on understanding the spatial distribution of discrete events (i.e., cells) by comparing observed patterns to a null hypothesis of Complete Spatial Randomness (CSR). Under CSR, point events are assumed to occur independently and uniformly across space. Deviations from CSR generally arise due to two fundamental types of spatial effects, first-order effects and second-order effects (26).

First-order effects reflect large-scale spatial variations in point intensity driven by environmental influences or tissue architecture. These are typically manifested in differences in point abundance across the landscape. Quantification of first-order effects includes summary metrics such as the total number of points, class-specific counts and densities, and the proportional composition of different point classes.

Second-order effects arise from interactions among individual points and influence the fine-scale structure of the spatial pattern. These effects are assessed using second-order summary statistics that evaluate spatial relationships between pairs of points, thereby revealing biotic interactions occurring at local scales. Key second-order metrics include the Morisita-Horn Index, which measures compositional similarity between spatial populations; the Colocalization Index, adapted from ecological studies to quantify the spatial overlap between two point classes (18); the Nearest Neighbor Index, which evaluates proximity between different cell types; and Ripley’s H function, which assesses clustering or dispersion patterns across a range of spatial scales. Mathematical definitions and derivations for all second-order indices are provided in the **Supplementary Methods**.

### Regional-Level and Local Spatial Profiles

When spatial statistics are calculated locally, the result is a two-dimensional spatial map that captures variations in tissue architecture across a sample. These localized statistical profiles are particularly valuable for investigating spatial heterogeneity, enabling the identification of microenvironmental differences that may not be apparent from global analyses.

Local density maps represent spatial profiles of first-order indices, including local cell densities estimated using kernel smoothing techniques such as Gaussian or Epanechnikov kernels (38,39). These maps can also incorporate additional spatial features, such as Euclidean distances from each location to tissue compartment boundaries or other macroscopic structures of biological relevance.

The Getis-Ord Gi*-score is another powerful local metric that detects statistically enriched or depleted regions for specific cell types (40–42), highlighting localized clustering or exclusion patterns within the tissue.

Furthermore, second-order point statistics—such as colocalization indices, nearest neighbor distances, and Ripley’s H function—can be adapted to local computation. In our methodology, these measures are estimated using spatial convolution smoothing to perform the necessary summations across neighborhoods. Specifically, we employ a Gaussian kernel to compute running values of each statistical index in a moving window across the tissue. This results in a high-resolution spatial profile in which each pixel is assigned a value representing the local neighborhood estimate of the index. Detailed mathematical formulations and implementation strategies for the convolutional approach are provided in the **Supplementary Methods**.

### Local Microenvironments (LME) Classification

While the optimal definition of tumor microenvironments should ideally incorporate biological insights including signaling pathways, expression profiles, and functional states, we propose an agnostic, data-driven framework to define Local Microenvironment (LME) categories using only cell-type abundances and spatial organization derived from point-location data.

This approach draws on methods from field ecology, where landscapes are divided into quadrats, small, spatially defined regions used to quantify local population distribution. To adapt this strategy, we apply a convolution bandwidth parameter that sets the spatial resolution for assessing each cell’s neighborhood. For each “quadrat”, we compute the abundance of each cell type and a mixing index, which quantifies spatial uniformity across “sub-quadrats” within that neighborhood. Instead of rectangular quadrats, we use circular regions defined by the Gaussian kernel implemented as the spatial convolution approach.

The mixing index is based on a univariate form of the Morisita-Horn index. It compares the observed cell distribution across sub-quadrats to a uniform (complete spatial randomness) expectation. A high mixing score indicates even spatial distribution, while a low score reflects clustering or segregation. Additional mathematical details of the mixing index computation and bandwidth kernel can be found in **Supplementary Methods**.

We define LME categories according to the combination of local cell densities and the mixing score. For each cell type, distributions of abundance and mixing score values are generated from all quadrat scores across all samples considered in a training set. This set should span the range of all expected microenvironments relevant to the study. Each of these distributions is then partitioned into three regions, representing low, medium, and high values. Because the mixing index is not well defined in the context of low-abundance distributions, we collapsed the nine original categories into three broader classifications that align with our LME framework. The criteria for this reclassification are as follows: the “B” (Bare) category denotes regions of low cell abundance, irrespective of the degree of mixing; the “S” (Segmented) category corresponds to moderate to high cell abundance with low mixing, indicating clustered distributions; and the “M” (Mixed) category represents moderate to high abundance with high mixing, indicative of more uniformly dispersed populations. Notably, due to the sensitivity of the mixing index to cell abundance, cases with medium abundance and intermediate mixing levels are also categorized as “Mixed” (M), since the index tends to underestimate mixing in the presence of low point counts.

Following this classification, each cell type within a defined quadrat is assigned one of the B, S, or M labels. A composite LME code is then generated by concatenating the labels across all tracked cell types within that region. In a system containing three distinct cell types, this results in 27 possible LME combinations (e.g., BBB, BSM, MMM), each representing a unique local ecological state (**Supplementary Methods**). To generate a spatially resolved map of the tissue landscape, we apply a convolution-based smoothing function, assigning each pixel the most probable LME category based on its surrounding microenvironment.

This approach yields a high-resolution classification of emergent spatial domains that are ecologically and biologically meaningful. It enables spatial stratification of tissue samples, identification of niche-level tumor features, and correlation with clinical phenotypes and treatment outcomes.

### Landscape Fragmentation and Composition Metrics

Tumor tissue can be conceptualized as an ecological mosaic—a spatial assembly of distinct biological niches or “patches” that correspond to non-overlapping regions defined by their cellular composition and microenvironmental context (43–46). To quantify the spatial complexity inherent in these tissue landscapes, we apply fragmentation analysis, a set of metrics designed to capture both the compositional and configurational properties of categorical tissue maps.

Composition refers to first-order spatial properties and quantifies the proportion of the tissue landscape occupied by each region type, such as tumor, stroma, or immune-dense areas. This provides insight into the relative abundance of different tissue compartments.

Configuration, by contrast, captures second-order spatial characteristics, describing how patches are spatially arranged, their geometric irregularity, and the degree to which they appear fragmented or aggregated within the overall tissue architecture.

This analytical framework is flexible and can be applied to a variety of spatial data sources, including regional-level spatial statistics, maps derived from Local Microenvironmental Ecology (LME) classification, or outputs from segmentation algorithms. The only requirement is that the input consists of a categorical spatial profile composed of discrete, labeled patches, enabling a reproducible and biologically interpretable analysis of tissue organization.

Landscape metrics used to characterize tissue architecture can be broadly categorized into three major classes: geometrical properties, shape metrics, and diversity metrics.

Geometrical properties describe the fundamental physical attributes of spatial patches, such as area, perimeter, and the total edge length at interfaces between adjacent regions. These metrics also include normalized area-to-perimeter ratios. Together, they provide insight into class dominance, rarity, and the extent of potential interfaces for cell-cell or region-region interactions, which may influence tumor-immune dynamics or stromal boundaries.

Shape metrics assess the morphological complexity of individual patches. These include the perimeter-area ratio, which is sensitive to spatial scale; the shape index, which offers a scale-independent measure of shape complexity; and the fractal dimension, which captures deviations from idealized Euclidean forms and approximates contour irregularity. These measures help to quantify the structural complexity of regions within the tumor microenvironment.

Diversity metrics evaluate the variety and distribution of patch types across the tissue. Examples include the Shannon diversity index, patch density, the proportion of the landscape occupied by each class, and contagion, a measure of the degree of interspersion and adjacency between different patch types. Complete mathematical definitions and formulations for shape index, fractal dimension, and other derived metrics are provided in the **Supplementary Methods**.

Landscape fragmentation metrics can be assessed at three spatial scales: the patch, class, and landscape levels. At the class level, the patch-specific metrics are aggregated to summarize spatial attributes of each patch type. Key metrics include the total number of patches, cumulative area, and the proportion of the landscape that each class occupies. The patch density metric, the number of patches per unit area, serves as a general index of fragmentation, as higher densities often indicate greater disruption of contiguous regions. The largest patch index further reflects dominance by indicating the proportion of the landscape occupied by the most expansive patch of a given class. Similarly, edge density normalizes total boundary length to the landscape area, quantifying the extent of physical interfaces where cell-type mixing or functional transitions may occur.

At the landscape level, metrics such as the landscape shape index summarize edge complexity across all classes, while contagion, also referred to as interspersion, estimates the likelihood that neighboring pixels belong to the same class, thereby capturing spatial aggregation or dispersion of tissue elements. Lastly, the Shannon diversity index quantifies overall ecological diversity within the tissue by considering both the richness (number of distinct classes) and the evenness (distributional balance among them), offering a holistic view of tumor microenvironment heterogeneity. Metric equations, index formulations, and analytical thresholds are detailed in the **Supplementary Methods**.

Together, these hierarchical metrics provide a comprehensive lens for interpreting the spatial organization of tumor tissue. For example, a tissue with high patch density but small average patch area likely exhibits pronounced fragmentation, with frequent shifts in microenvironmental context. In contrast, a low contagion score would suggest spatial disorganization or fine-grained intermingling of distinct tissue regions, as might be seen in poorly differentiated tumors or areas of immune infiltration. Conversely, a high contagion or shape index might indicate well-demarcated regions or organized compartments, such as fibrotic stroma or tumor nests. Additionally, the Shannon index serves as a sensitive metric for spatial diversity, where high values reflect complex ecosystems with balanced tissue class distributions, while low values correspond to monocultures or highly dominant cell states. By synthesizing these spatial features, fragmentation analysis enables quantitative assessment of tissue architecture, identification of recurring spatial motifs, and potential correlation with clinical outcomes such as prognosis or treatment response.

### Data and Code Availability

All code for data processing and analysis associated with the current submission is available at the CarrLab GitHub repository: https://github.com/CarrLab/TLA.

All documentation for usage and future updates will be maintained in this web location.

## Results

A foundational approach to understanding spatial organization in biological tissues involves analyzing the distributions of discrete entities —such as cells or events— across space. In ecology, spatial point pattern analyses quantify the effects of population dynamics, niche partitioning, and inter-species interactions, which manifest as first-order effects (based on cell density) and second-order effects (based on spatial proximity) (25,26). These same principles apply to tumor biology, where spatial distributions of cells shape microenvironmental conditions that influence tumor progression, metastasis, treatment resistance, and adaptation.

In cancer, cellular spatial patterns are central to the formation of the TME and frame the evolutionary context of disease development. Local patterns of immune infiltration, stromal organization, and necrosis mirror ecological systems, linking single-cell phenotypes to tissue-level functionality. Cellular organization governs signaling, apoptosis, proliferation, and differentiation, and disruptions in these configurations can drive pathological remodeling (27). Viewing tumors as ecological systems motivates the integration of landscape ecology and spatial statistics into digital pathology pipelines (28–31).

To apply landscape statistics to a tissue, the cells first must be identified and classified into different cell types (generally through digital pathology techniques). Then, landscape ecology statistics can be applied to the tissues at four spatial scales: the cell level, the regional/local level, local microenvironments, and landscape fragmentation.

We present results for examples that illustrate outcomes for the three data formats introduced in the methods section. In each case, we show some typical features quantified using our methodology.

### Whole-Cell Detection

First, whole-cell detection methods use highly multiplexed imaging to identify and phenotype individual cells across a tissue section. For example, Keren et al. (2018) applied multiplexed ion beam imaging (MIBI) to analyze triple-negative breast cancer samples using a 36-marker panel (22). A convolutional neural network (CNN) was trained to classify cells based on marker expression, producing cell-level phenotype maps with corresponding spatial coordinates. The output for each sample was a rasterized map of individually segmented cells annotated with phenotypic labels, as shown in **Figure 2**.

**Figure 2.**
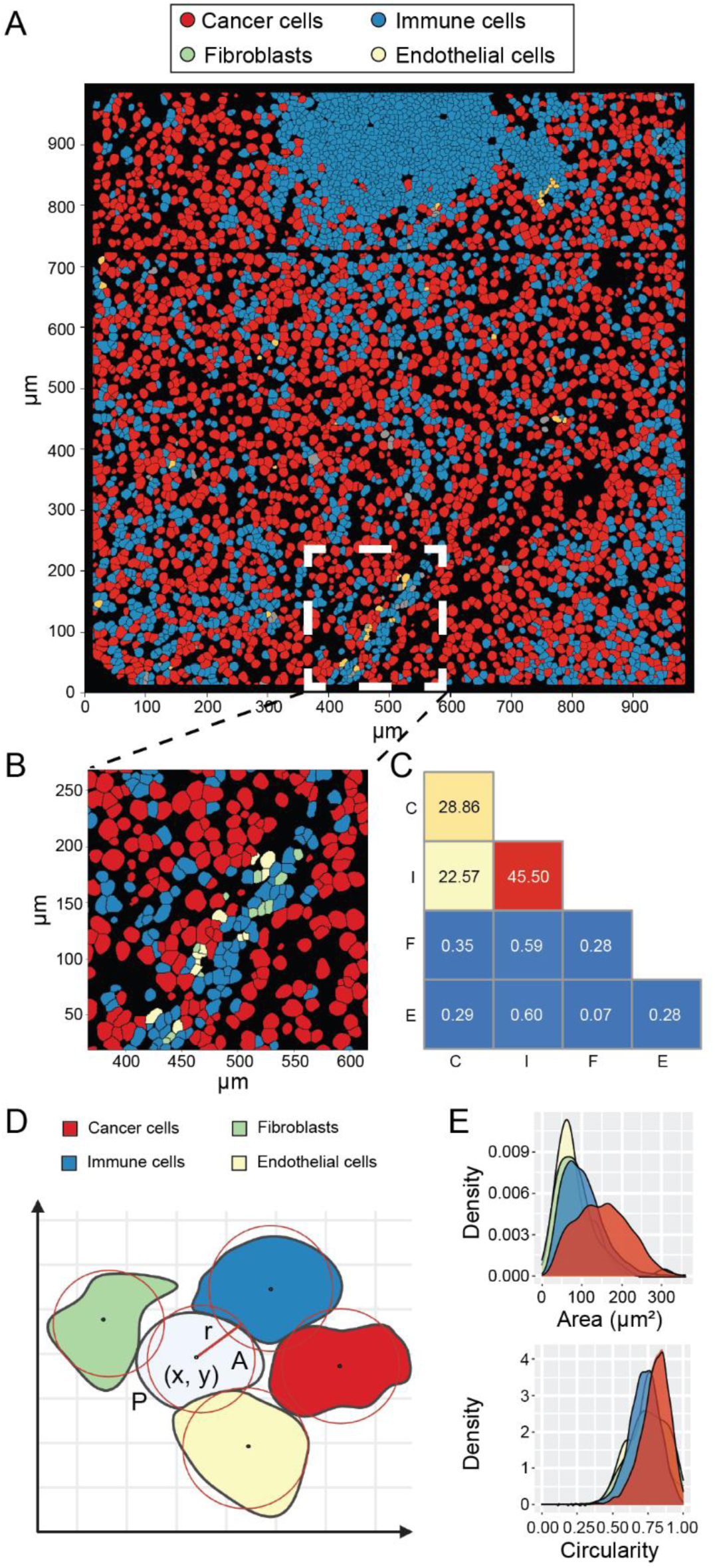
Whole-cell Detection Using Multiplexed Ion Beam Imaging of Triple Negative Breast Cancer. A. Example of a raster map of individually identified whole cells, colored by phenotype. The whole sample for this methodology is only 1mm x 1mm. B. Small 200um x 200um section showing details of cell contacts. The difference in sizes between cell types is evident. C. Heatmap representing frequencies of adjacency contacts between cancer cells (C), immune cells (I), Fibroblasts (F) and endothelial cells (E). Adjacencies of immune cells with other immune cells correspond to 45.50% of the interactions in this sample, while cancer cells with cancer cells are 28.86% of them, and cancer cells with immune cells are 22.57%. All the rest of the interaction pairings are rare (<1%) (indicated by blue). D. Diagram showing some of the morphological properties calculated for individual cells, including area A, perimeter P, equivalent radius r and centroid (x, y). E. Distributions of cell area and circularity for each cell type class. The circularity is calculated from the area and perimeter as C= “exp” (1-P^2^/4Aπ), which equals 1 for a circular shape and goes down to 0 for less circular shapes, elongated or very dendritic. In this sample, cancer cells are indeed typically larger in area and more circular than other cell types.

This data is used to make measurements for basic cell morphology, including area and circularity, as well as cellular context accounted by the frequencies of adjacencies. More statistics can be collected for this level of analysis, including other cell morphology features, more sophisticated measures of context that weight adjacencies by the edge length of each contact, and accounting of expression profiles at single-cell levels, given that multiplex platforms provide such richness of data. Additionally, centroid coordinates can be estimated geometrically for each cell and used downstream for point statistics, LME classification, and fragmentation analysis. However, our examples of these analyses use other data sources mainly because the MIBI platform is limited in the size of the mapped landscape. These samples are only 1mm across, which constrains the capacity to detect large-scale tissue features and long-range effects. Also, some of the metrics implemented in these analyses require segmenting the region of interest (ROI) into smaller quadrats to build spatial distributions, and each of these quadrats needs to contain an appropriate number of cells to grant statistical robustness; this is not feasible with such a small field of view. Thus, we limit the MIBI analyses to a few cell-level statistics.

In **Figure 2** we show a sample case. Individually identified cells are mapped out and colored by cell type (A and detail in B). We show examples of the distributions of cell areas (D) and circularity (E) for this slide, as well as a heatmap diagram showing the frequencies of adjacencies (C). These results demonstrate that tumor cells in this TNBC sample are consistently larger and more circular than other cell types. Also, lymphocytes tend to be in contact preferentially with other lymphocytes (as is evident by the large cluster of lymphocytes in the lower side of the sample), and tumor cells also preferentially interact with other tumor cells, as well as lymphocytes as a close second.

### Cell Location Data

Nuclear segmentation has been used to characterize cell types in hematoxylin and eosin (H&E)-stained histology. In studies of ductal carcinoma in situ (DCIS), CNN-based classifiers were trained to assign nuclear centroids to specific cell phenotypes (34,35). These analyses identified three major cell populations: epithelial cells, immune cells, and other stromal cells. The spatial format for each biopsy comprised a map of labeled nuclear centroids, with each point denoting the location and phenotype of an individual nucleus (**Figure 3**).

**Figure 3.**
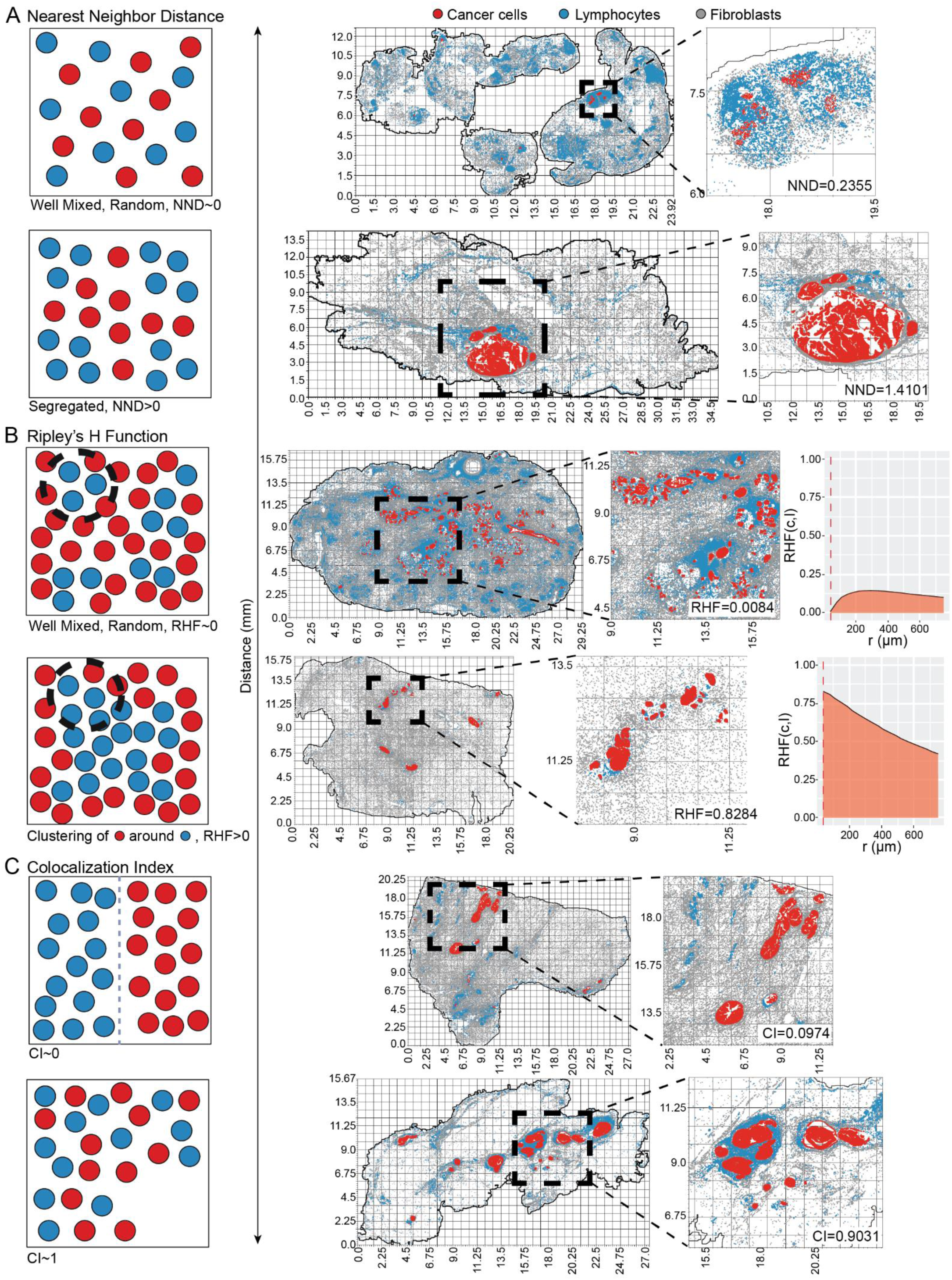
Cell-Level Spatial Metrics in DCIS. A. Examples for extreme cases of the Nearest Neighbor Distance index. The top panel row displays a diagram of a well-mixed situation, with NND close to 0, showing that different cell types are typically uniformly distributed with respect to each other. To the right there is a real data example showing the NND of lymphocytes (blue) around tumor cells (red) averaged over tumor cells is low, indicating that lymphocytes are typically intermixed with tumor cells. The second row shows a diagram of a segregated state, in which the NND between red and blue dots are long. To the right a real data example of this situation is presented, in which the separation between tumor cells (red) and lymphocytes (blue) is evident. Both examples correspond to the NND of lymphocytes around tumor cells, which actual value of the index shown in the lower right corners. B. Examples for the Ripley’s H index are shown. This is a metric calculated for a specific scale, d=40um. In this case, the separation being quantified is between features of this specific scale size. Thus, the mixed case, upper left diagram, shows features well mixed together at the resolution scale of the circle. The lower left diagram shows the separation between the point classes at that same scale. In real data examples, in which we can see groups of tumor cells (red) and groups of lymphocytes (blue) intermixed together (upper right) and separation between clusters regardless of the scale (lower right). Actual index values are shown in the inset box. Curves of the value of the index for larger scale values are shown to the far right in both cases, showing that in the first example the index increases slightly with d and then drops after a scale of 200um, while in the lower example it consistently drops, but not a lot. It is generally expected to observe lower values at longer scales as small features get smoothed out at such scales. C. Examples for the Colocalization Index, which quantifies the similarity between two spatial distributions of points. In the upper case, the diagram shows two segregated distributions of points, with a value of the index close to zero, which can be observed in the real example on the right-hand side. Whereas the lower example shows a situation in which the two distributions are like each other, and an index close to one. The real case example for this case is not obvious. Similarly to the RHF index, there is a set scale for the calculation of this index, in this case given by the size of the bins used to construct a discrete spatial distribution (i.e. as a histogram). The scale used for these examples is 40 μm. Even though lymphocytes (blue) are not exactly infiltrated in clusters of tumor cells (red), at the scale of smoothing the regions where there is a high density of tumor cells also have a higher density of lymphocytes. This example is important to indicate a caveat in using this metric as a measure of infiltration.

Many measurements can be derived from a point-statistics analysis. This analysis will be published in much greater detail elsewhere, but here we present key examples to illustrate our methodology and to display the behavior of different point statistics measures. Additionally, this type of data can be used downstream for creating LME categories and fragmentation analysis. To keep this article concise, we limit these results to examples of spatial statistics metrics and the LME construction (**Figure 4**).

**Figure 4.**
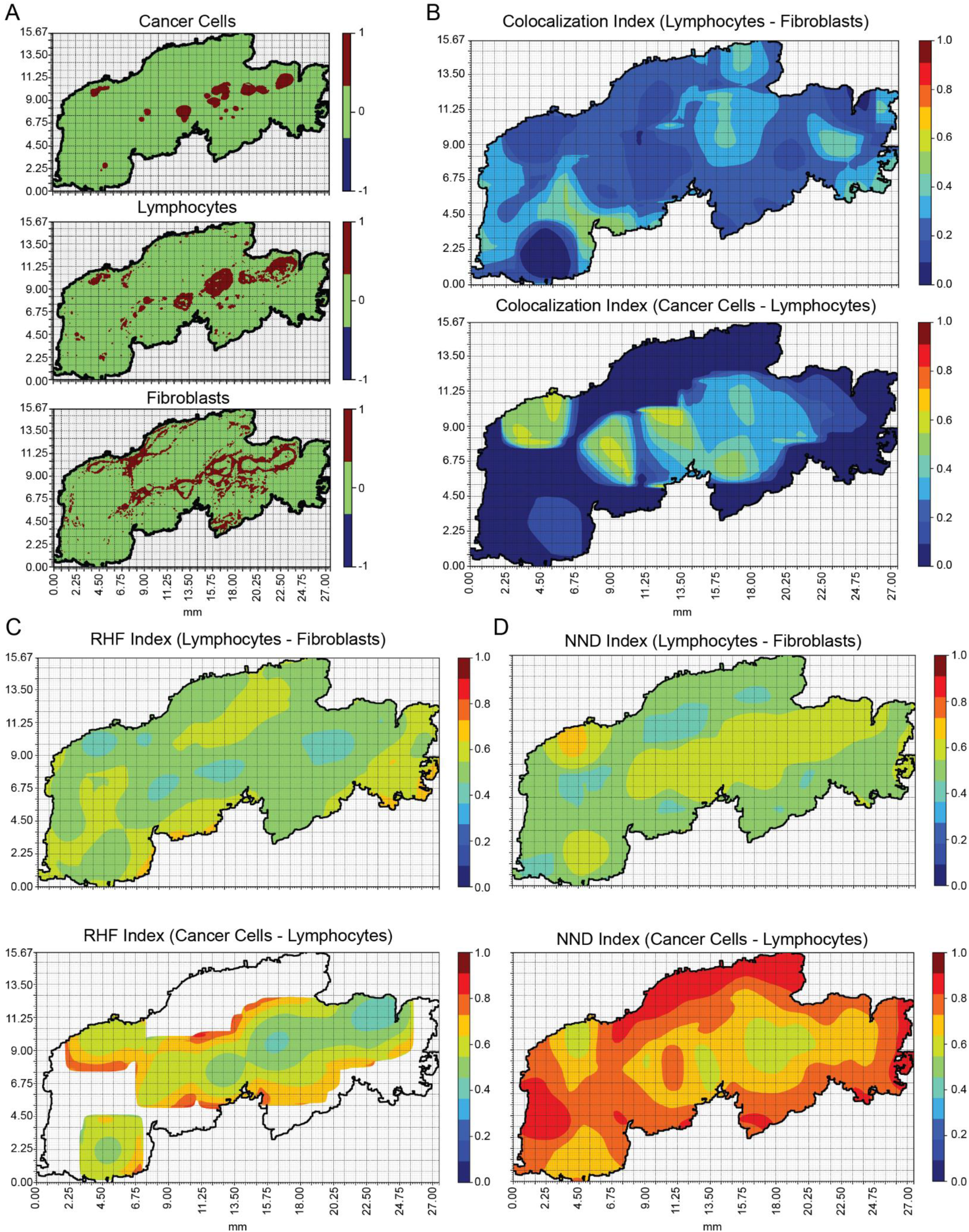
Regional-Level and Local Spatial Profiles in DCIS. A. Hot and cold density regions estimated using the Getis-Ord Gi* statistic, for each of the cell phenotypes in the DCIS study. Regions colored red are hot spots where the density of cells is larger than expected according to the total cell density of the sample (i.e. statistical significance using a z-score test). Green indicates regions where the cell density is within the range of expected values. Blue regions (not observed here) indicate cold spots where cell density is lower than expected. B. Local Spatial Profiles for the Colocalization Index are shown for two examples, between lymphocytes and fibroblasts and between cancer cells and lymphocytes. The colocalization index is calculated in the neighborhood around each pixel using a spatial convolution with a Gaussian kernel with a 20um bandwidth. In this way, these profiles represent local variations of these metrics, in this case showing that most of the slide has moderate to low values of colocalization, with some small regions where the distributions of cell types are similar. C. Local Spatial Profiles for the Ripley’s H function of fibroblasts around lymphocytes and lymphocytes around tumor cells, calculated locally using a spatial convolution method with a Gaussian kernel with a 20um bandwidth. The extent of the slide shows mostly intermediate values, with some larger values in the periphery of the middle region, where tumor clusters are, indicating that segregation of cell types happens away from those areas. D. Local Spatial Profiles for the Nearest Neighbor Distance index of fibroblasts around lymphocytes and lymphocytes around tumor cells, calculated locally using a spatial convolution method with a Gaussian kernel with a 20um bandwidth. Like panel C, larger values indicate segregation of cell types whereas small values indicate closer proximity between cell types.

In **Figure 3** we can see examples for each of the point-statistics metrics, a schematic that explains extreme cases for each measure, and centroid overlays of specific real cases that represent each of these extremes. Nearest-neighbor Distance Index (NND) (**Figure 3A**): the diagrams show that a low value of this index is a signature of CSR, meaning that the two cell types considered (in these cases they are DCIS cells and lymphocytes) are well mixed and the frequency of finding a lymphocyte next to a DCIS cell is like the frequency of finding another DCIS cell. This effect is evident in the detail shown on the right-hand panel, where we can see that lymphocytes are thoroughly mixed and infiltrated in the clouds of DCIS cells. On the other hand, large values of NND indicate that the typical distance from DCIS cells to the closest lymphocytes is larger than the distance to the closest DCIS cell, hence representing a case of segregation of lymphocytes away from DCIS cells. This can be seen in the right-side detail panel where lymphocytes are excluded from the DCIS cell clusters.

Ripley’s H function (RHF) (**Figure 3B**): in this case again, a low extreme is indicative of CSR. An important distinction is that the properties conveyed by this metric have a characteristic scale defined by the radius used to calculate *H*. A way to understand this metric is similar to the principle for NND, but instead of considering just the first neighbor, count the ratio of the number of points of each type inside a circle of radius *d*. Therefore the “mixing” or “clustering” is defined with respect to this scale. For this particular example *H*was calculated at a scale of *d*=40μ*m*(a small green circle in the lower left corner shows the dimension of the scale used). In this sense, cells could be clustered forming features, and this metric could still have a low value if those features are smaller than the scale *d*and they, as coarse-graining features instead of the individual cells, are well mixed and satisfy a CSR condition. We can see this in our example, on the right-hand side we see clusters of DCIS cells mixed with lymphocytes providing a well-mixed condition at the scale used, though at the cell scale, this example would show separation between the two populations. In contrast, the other extreme case clearly shows a separation between the populations indicating a typically large distance from DCIS cells and groups of lymphocytes.

Colocalization Index (MHI) (**Figure 3C**): for the case of the colocalization index, measured employing the Morisita-Horn index. The high extreme (*M* ∼ 1) is not necessarily a case of CSR, though CSR would certainly have this result. In a more general sense, what this metric indicates is the similarity between the spatial distributions. If the spatial distribution of the two cell types is not uniform, but are similar to each other, we will observe this extreme, as illustrated in the lower panel. The scale of the quadrats used, 750×750[μ*m*^2^], comes into play in much the same way as the scale for RHF. As we see in the real data example, lymphocytes follow the spatial distribution of DCIS cells, suggested by the fact that regions with large concentrations of DCIS cells also have larger concentrations of lymphocytes, even though at lower scales they are not infiltrated. On the other hand, the other extreme *M* ∼ 0 reflects a condition of total separation of the two populations, at the scale of the spatial distribution binning, as is clearly shown in the example case.

### Region Detection Data

We present region-level results from two studies. The first one is comprised of regional results derived from the point-statistics analysis in our DCIS example (see previous section). In **Figure 4** we show the spatial profiles for local measurements of NND, RHI, and MHI, showing how these indices can be estimated in small regions to represent the variations in space.

For the spatial profile of NND of DCIS cells around lymphocytes, we can see a region of negative values, whereas in most other regions we see values typically close to zero. A negative value of this metric indicates that DCIS cells (test cells) are closer to any lymphocytes (reference cells) than other lymphocytes, this happens when the density of reference cells is small while that of test cells is large, meaning that test cells are common in the region and reference cells are individually infiltrated and surrounded by test cells. On the other hand, a positive score indicates clustering of test cells around reference cells, while close to zero values indicate an even distribution. In this sample, the distributions of DCIS cells and lymphocytes are pretty even across the tissue landscape. For the local Ripley’s H score, we show an example of lymphocytes around other lymphocytes that show regions of clustering of these types of cells. Notice that the reference cell density is calculated locally, thus the local values of H reflect clustering within regions (which is a circle with *r* = 0.75mm). This profile is thus a display of the intensity of local relative variations of the density of lymphocytes. Finally, for the colocalization index, we present an example of the overlap of DCIS cells and lymphocytes, showing that for the most part, the distributions of these cell types are very similar across the sample, with some variations in regions on the edges of it that display moderate separation of the distribution patterns, mostly driven by low densities of lymphocytes.

In addition to the three basic point statistic metrics, we also estimated the profile for the Getis-Ord Gi* statistic for the density of lymphocytes. This is a regional statistic that detects locations in which there is higher, or lower than expected cell density for a particular cell type. We show specifically the regions that are statistically significant for the z-statistic, corresponding to hot (red) and cold (blue) regions for cell density, while green regions indicate expected densities. Interestingly, this sample shows quite a lot of local clustering across the sample, including a middle region where DCIS cells are concentrated.

Finally, in landscape ecology, ecosystems are often decomposed into spatially distinct biological niches, referred to as “ecologic phenotypes” or LME, which represent specific combinations of species coexisting within environmental contexts. Analogously, tumor tissues can be understood as ecological mosaics composed of functionally distinct regions, each shaped by local cellular composition and spatial configuration. We present an example of the spatial profile of LME categories defined by the local abundance and mixing scores for each cell type, as described in the methods (**Figure 5**). We can see that this strategy is capable of differentiating regions with different population configurations as different types of micro-ecologies. There is a main region with profiles for all DCIS cells that are both abundant and mixed with segregated lymphocytes (red region) and a region with segregated DCIS cells and lymphocytes (green area) (**Figure 5B**). This separation is evident from the point distribution profiles separated by classes (**Figure 5C**).

**Figure 5.**
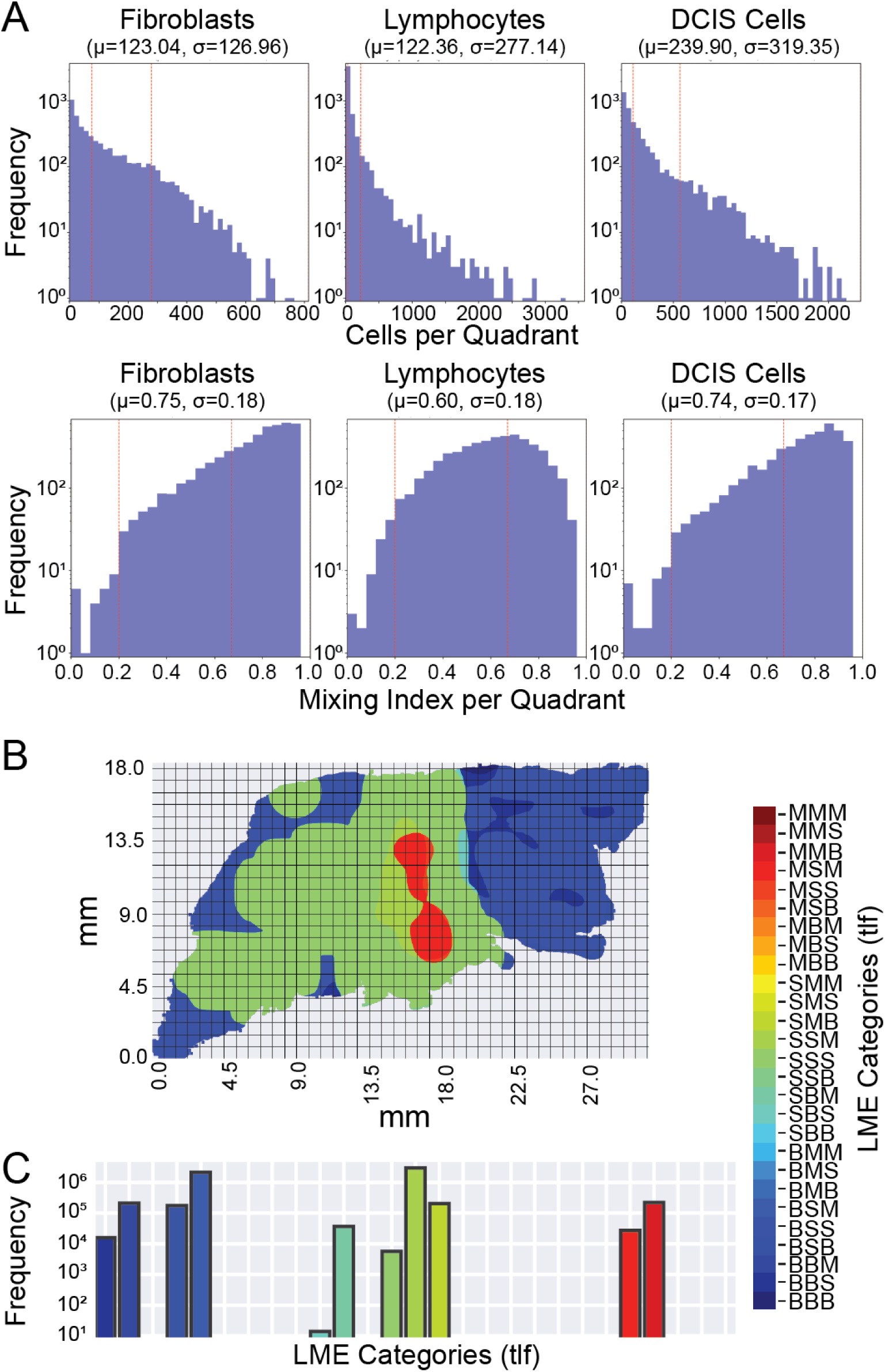
Local Microenvironments (LME). A. Local values of cell abundance and mixing for all samples in the study are collected and summarized. Distributions of values are used to assign low, medium and high levels, as observed collectively in all samples. B. Example of LME classification based on local values of cell abundance and mixing. C. Quantified frequency of LME subtypes of the example landscape in panel B. (S) Segmented environments are those in which cells are clustered together; they have moderate to high abundance and low mixing. (M) Mixed environments are those where cells are mixed uniformly; they have moderate to high abundance and high mixing. Because the mixing score is sensitive to low abundance, medium abundance levels are considered “mixed” (M) if the mixing is medium, as mixing is biased to lower values with lower abundances. With its scheme, (B, S, M) codes are assigned for each individual class, forming categories like: ‘BBB, BBS, BBM …. MMM’. Each of these categories represents a basic type of Local Micro-Environment that is consistent across the entire study and can be used to define ecological phenotypes in the TLA.

As an unsupervised and quantitative framework, LME analysis provides a reproducible and scalable method for partitioning tumor microenvironments using only spatially resolved cell type data. While it is not intended to replace biologically curated models, this strategy offers a principled way to delineate tumor architecture and immune dynamics, especially when integrated with molecular profiling or clinical annotation.

All these spatial profiles can be further analyzed using fragmentation analysis tools to measure the morphological and configuration properties of the “habitats” in the sample. For this, it is necessary to convert them into discrete maps composed of a set of patches or local habitats. This is already the case for the Getis-Ord hot/cold region profiles and the LME profiles. In the case of the local statistics profiles, a possibility is to stratify the continuous values into discrete contour levels and use those as the patches for the fragmentation analysis. In this sense, if we visualize these profiles as surface landscapes with mountains and valleys, the fragmentation analysis of contour levels is a way to assess the ruggedness or smoothness of such a surface.

### Fragmentation Statistics

Our last example is the regional classification data of PDAC resections. In this case, discrete regions representing different tissue compartments are identified using machine-learning image processing. The regional segmentation data does not allow for point-statistics analysis, but it can be processed using our fragmentation analysis approach. In **Figure 6** we show some examples of fragmentation metrics that were significant in this study. Here we present extreme example profiles for the Shape Index, the Patch Density, and Contagion metrics. In this study, these metrics were predictive of the time to recurrence after neoadjuvant therapy in PDAC. The shape index is a normalized measure of the perimeter-to-area ratio of the patches conforming to the landscape. A large value of this index indicates that the typical patches have irregular and complex shapes which can be an indication of fragmentation and suggests a dynamic in which signaling and other processes of transport or exchange of chemical substrates between niches. We see that the lower extreme case shows typically small patches with globular and compact shapes, while the higher extreme case shows elongated and complex-shape patches. The nature of interactions between patches would be radically different in these two cases. The patch density is simply a measure of the number of patches per unit area. The extreme cases are clear, showing a case with a moderate number of patches, especially stroma, for the lower extreme and a very rich and populated landscape for the higher extreme, with many small tumor patches and a more fragmented stroma. Finally, contagion is a metric that quantifies the level of cohesiveness of the landscape, expressed as the probability that two randomly picked neighboring locations are of the same class.

**Figure 6.**
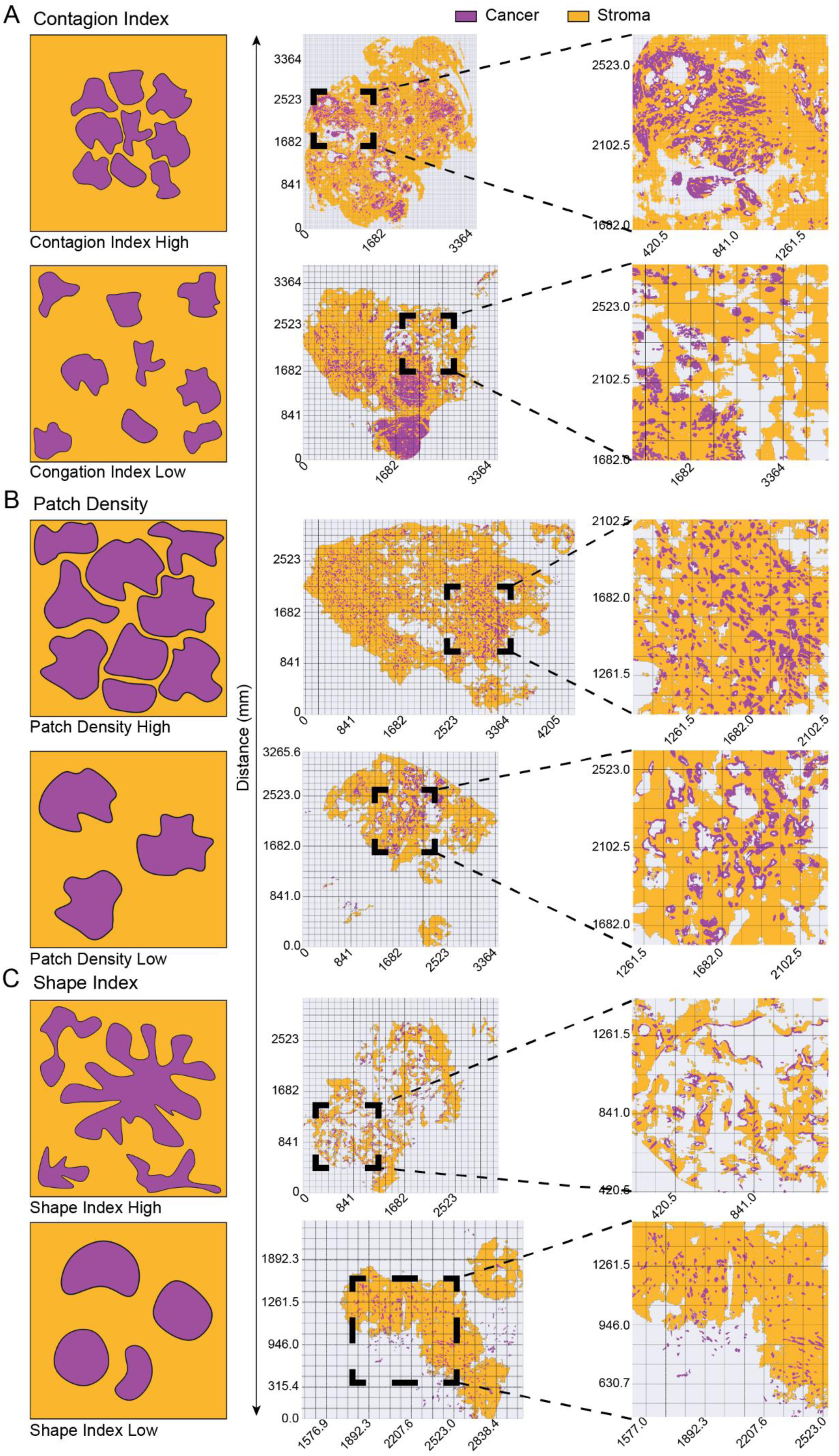
Landscape Fragmentation and Composition Metrics in PDAC. A. The Contagion Index: this is a metric of diversity and complexity of a landscape mosaic that quantifies the probability that two randomly chosen neighboring locations (pixels) belong to the same patch class, thus measuring the level of connectivity, or conversely fragmentation, of the landscape. These are two case examples showing extreme high (upper case) and extreme low (lower case) of this metric in the PDAC study data, and explanation diagrams (right hand panels). B. The patch density: the number of patches, of any class, per unit area. Two examples and explanation diagrams are shown. C. The shape index: this is a morphology measure that accounts for the relation of perimeter and area that is adjusted for size. Since the perimeter of a patch scales linearly with its length scale, while the area scales as the square of it, this measure is defined as a fourth of the perimeter divided by the square root of the area and is a measure of shape independent of scale. Compact, circular or squared, shapes have low perimeter to area ratio, and thus have low shape index values, while more complex, or dendritic, shapes have longer perimeters for the same area, thus having large shape index. Examples and diagrams of extreme cases are shown.

In the lower extreme case, we could see that depending on the location in the landscape, the two types of regions can be very intertwined, especially in the middle of the sample, suggesting that it’s generally likely that two random neighboring locations would be of different classes. The opposite is true for the other extreme case. In which, though the tumor stroma is very segmented, the stroma is much more prevalent in space and two random neighboring locations will likely be stroma. It’s important to point out that this measure is normalized for the abundance of each class, but a high value of contagion indicates the likelihood of two neighbors being of the same class, but it says nothing about the balance of the probabilities of different classes.

Third, region detection approaches segment larger structural compartments of the tissue. In pancreatic ductal adenocarcinoma (PDAC), a machine learning model was used to delineate tumor and stromal regions in post-treatment surgical resection samples (36). This region-level segmentation enabled the generation of spatial maps that facilitate downstream analyses, including the identification of specific cell types within distinct tissue compartments and investigations of spatial relationships, such as proximity to the basement membrane or blood vessels (**Figure 6**).

## Discussion

Interactions between different cell types in the TME shape disease progression and influence therapeutic outcomes. Those interactions and the ecology of the TME can be studied through spatial analyses of the different cell types. While technological advances in spatial biology have enabled increasingly detailed characterizations of tissue architecture, many current analytic approaches still rely heavily on reductionist models or narrowly focused metrics. In this study, we present a comprehensive analytical framework, the Tumor Landscape Analysis (TLA) pipeline, that adapts core principles from landscape ecology to better capture the complexity of tumor ecosystems. This framework moves beyond conventional spatial biology by offering a multiscale, quantitative, and ecologically informed view of the TME.

Conventional spatial analyses typically depend on identifying specific cell types and evaluating their relationships through simple distance metrics, adjacency maps, or nearest-neighbor interactions. These methods, while informative, often operate in isolation from the broader context of the tissue’s spatial organization. They rarely account for higher-order spatial properties such as fragmentation, aggregation, or habitat structure—features that are central to ecological systems and, by extension, highly relevant to tumor biology. The TLA framework addresses these limitations by incorporating a broader spectrum of spatial statistics, adapted from ecological theory. Metrics such as the Morisita-Horn index, Ripley’s H function, and the Shannon diversity index allow for the characterization of tissue landscapes in terms of both cellular composition and spatial configuration. Rather than focusing exclusively on pairwise cell interactions or compartmental delineations, TLA provides a means to assess how cells and regions are distributed, how they interact across distance scales, and how these patterns may reflect underlying biological processes such as immune infiltration, stromal remodeling, or clonal expansion.

One of the key innovations introduced here is the concept of Local Microenvironments (LMEs), defined not by prior biological assumptions but by empirical patterns of cell abundance and spatial uniformity. This unsupervised classification yields discrete ecological niches within the tissue, offering a reproducible and interpretable means of partitioning tumor architecture. Importantly, this approach is data-driven yet biologically intuitive: regions identified as “segmented,” “mixed,” or “bare” correspond closely with phenomena observed in pathology, such as lymphocyte clustering, immune-excluded zones, or necrotic cores. In parallel, the application of fragmentation metrics, long used in landscape ecology to describe habitat loss and environmental disruption, provides a novel lens through which to assess tumor morphology. Measures such as patch density, shape complexity, and interspersion offer insight into the degree of spatial disorganization within a tissue. These attributes are not merely descriptive; they have the potential to serve as spatial biomarkers, capturing aspects of tumor behavior that may correlate with aggressiveness, treatment resistance, or recurrence risk. They are likely to change dramatically with treatment.

Perhaps most significantly, TLA allows for integration across data types and spatial scales. Whether analyzing whole-cell segmentations, centroid point maps, or region-level classifications, the framework applies a consistent set of spatial tools that enable comparisons across tissue types, disease states, and imaging platforms. This adaptability is essential for translating spatial biology into clinical practice, where tissue samples vary in quality, modality, and resolution. Looking ahead, the potential applications of this framework are substantial. When applied to longitudinal or post-treatment samples, TLA could facilitate the reconstruction of tumor evolution under therapeutic pressure, revealing how spatial features emerge, shift, or regress. Its integration with molecular or transcriptomic data could yield composite ecological-genomic signatures, providing a richer context for understanding tumor behavior and cell-cell interactions. Moreover, TLA-derived metrics could be incorporated into predictive models or decision-support tools aimed at guiding therapy selection or patient stratification.

Taken together, the Tumor Landscape Analysis framework represents a shift in how we conceptualize and quantify tumor ecosystems. By drawing on principles from landscape ecology, it enables a more holistic understanding of spatial heterogeneity, one that captures not only what cell types are present in the TME, but how they are organized, how they interact, and what patterns emerge from their collective behavior. This perspective is both timely and necessary, as oncology continues to move toward more personalized, spatially informed models of care.

## Acknowledgments

This work was also supported by the Gerstner Family Foundation Career Development Award, the Grand Forks Career Development Award, and the Mayo Clinic Center for Clinical and Translational Science (CCaTS) through a Small Grant (UL1TR000135). Ryan Carr gratefully acknowledges this funding support, which was instrumental in the development and execution of this study.

## Notes

**Conflict of Interest Statement:** The authors declare no conflicts of interest.

### Competing Interest Statement

The authors have declared no competing interest.

### Summary of Updates

Minor writing edits and re-ordering of figures.

