## Supplementary Methods for "Tumor Landscape Analysis: An Ecologically Informed Framework to Understand Tumor Microenvironments"

***Equivalent radius****: if*
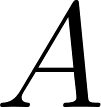
<https://www.codecogs.com/eqnedit.php?latex=A#0> *is the area of a spatial feature (e.g. a cell) then the equivalent radius*
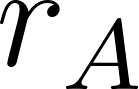
<https://www.codecogs.com/eqnedit.php?latex=r_A#0> *is the radius of a circle with area*
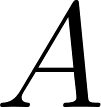
<https://www.codecogs.com/eqnedit.php?latex=A#0>*, and thus:*

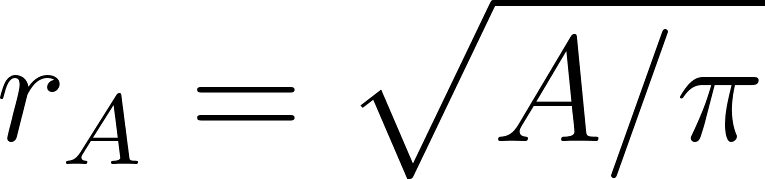
<https://www.codecogs.com/eqnedit.php?latex=r_A%3D%5Csqrt%7BA%2F%5Cpi%7D#0>

***Circularity measure****: if*
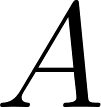
<https://www.codecogs.com/eqnedit.php?latex=A#0> *is the area of a spatial feature (e.g. a cell) and*
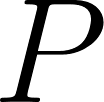
<https://www.codecogs.com/eqnedit.php?latex=P#0> *is its perimeter, then the area of a circle with perimeter*
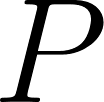
<https://www.codecogs.com/eqnedit.php?latex=P#0> *is*

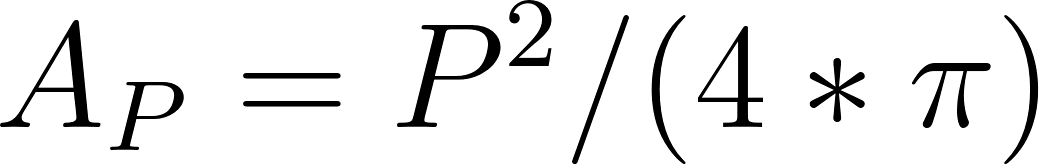
<https://www.codecogs.com/eqnedit.php?latex=A_P%3DP%5E2%2F(4*%5Cpi)#0>

*In the plane, a circle encloses the largest area for a given perimeter, so*
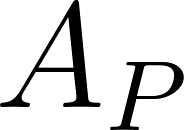
<https://www.codecogs.com/eqnedit.php?latex=A_P#0> *is a maximum for a given*
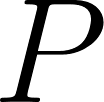
<https://www.codecogs.com/eqnedit.php?latex=P#0> *and we can define the excess area of any arbitrary shape as:*

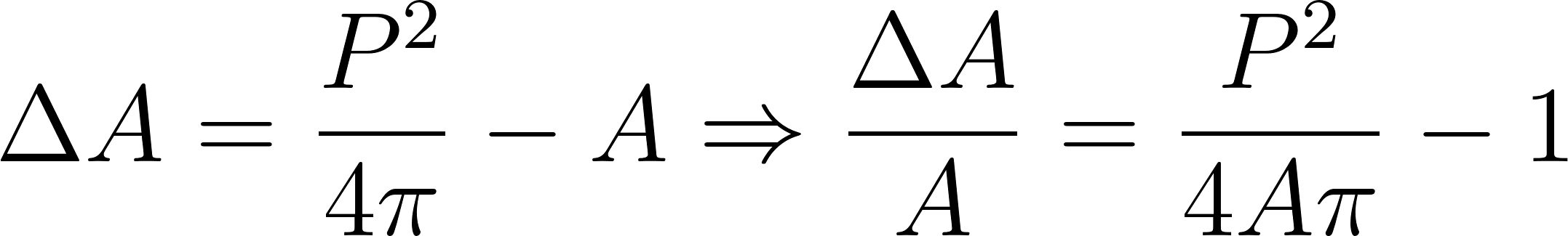
<https://www.codecogs.com/eqnedit.php?latex=%5CDelta%20A%20%3D%20%5Cfrac%7BP%5E2%7D%7B4%5Cpi%7D%20-%20A%20%5CRightarrow%20%5Cfrac%7B%5CDelta%20A%7D%7BA%7D%20%3D%20%5Cfrac%7BP%5E2%7D%7B4A%5Cpi%7D%20-%201%20#0>

*And finally, we can define the circularity of any shape with area*
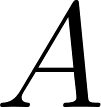
<https://www.codecogs.com/eqnedit.php?latex=A#0> *and perimeter*
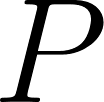
<https://www.codecogs.com/eqnedit.php?latex=P#0> *as the quantity:*

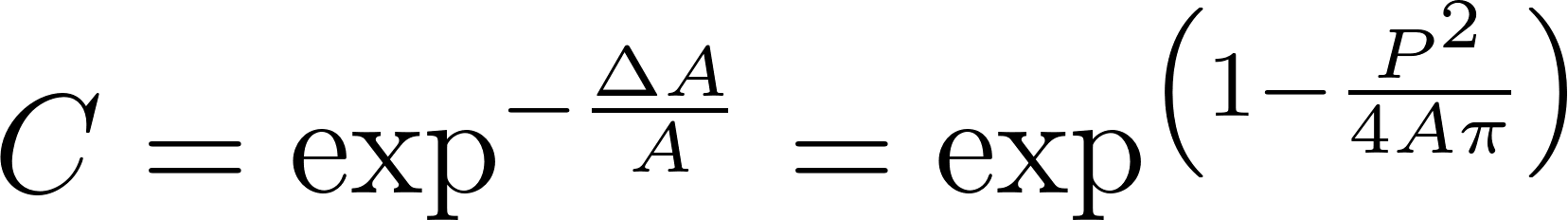
<https://www.codecogs.com/eqnedit.php?latex=%20C%20%3D%20%5Cexp%5E%7B-%5Cfrac%7B%5CDelta%20A%7D%7BA%7D%7D%20%3D%20%20%5Cexp%5E%7B%20%5Cleft(1%20-%20%5Cfrac%7BP%5E2%7D%7B4A%5Cpi%7D%5Cright)%7D#0>

*With*
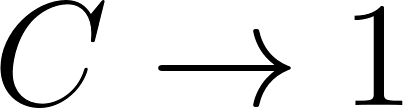
<https://www.codecogs.com/eqnedit.php?latex=C%5Crightarrow%201#0> *if the shape is close to a circle (i.e.*
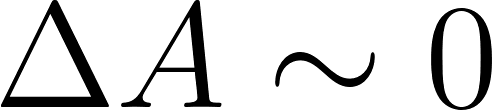
<https://www.codecogs.com/eqnedit.php?latex=%5CDelta%20A%20%5Csim%200#0>*) and*
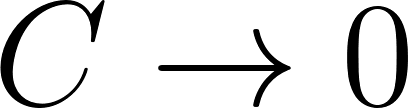
<https://www.codecogs.com/eqnedit.php?latex=C%5Crightarrow%200#0> *if the shape has a large excess area, i.e. an elongated or irregular amoeboid shape.*

***Number of neighbors:*** *if*
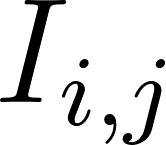
<https://www.codecogs.com/eqnedit.php?latex=I_%7Bi%2Cj%7D#0> *is the Indicator Function (equal to 1 if*
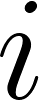
<https://www.codecogs.com/eqnedit.php?latex=i#0> *and*
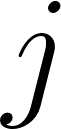
<https://www.codecogs.com/eqnedit.php?latex=j#0> *are adjacent neighbors and 0 otherwise) then the number of cells in contact with a cell*
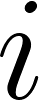
<https://www.codecogs.com/eqnedit.php?latex=i#0> *is*

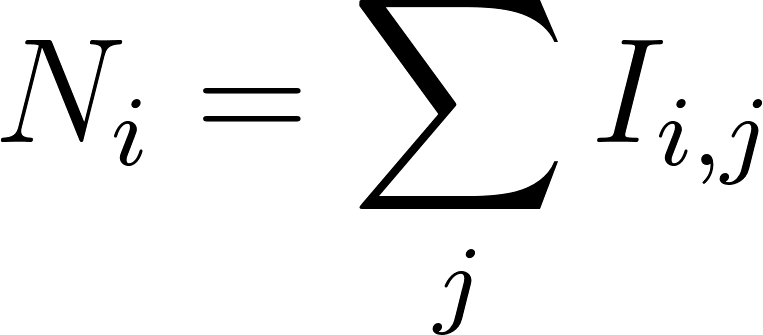
<https://www.codecogs.com/eqnedit.php?latex=N_i%20%3D%20%5Csum_j%20I_%7Bi%2Cj%7D#0>

*And the fraction of the number of neighbors of*
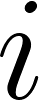
<https://www.codecogs.com/eqnedit.php?latex=i#0> *that corresponds to a particular cell class 𝛂.*

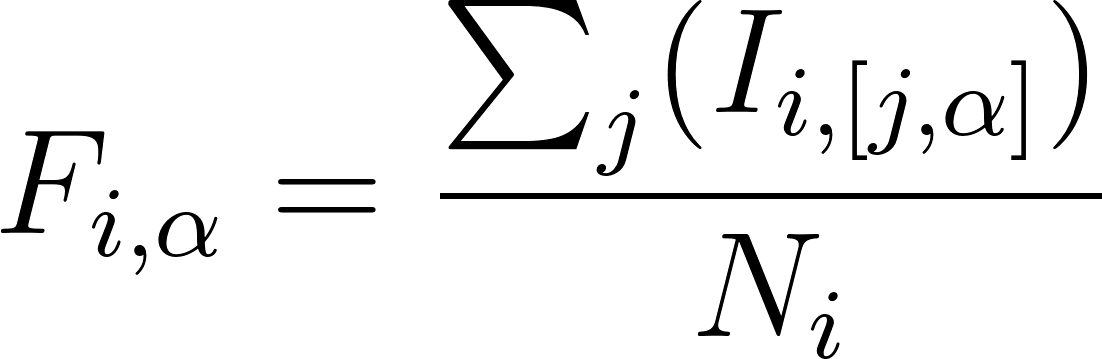
<https://www.codecogs.com/eqnedit.php?latex=F_%7Bi%2C%5Calpha%7D%20%3D%20%20%5Cfrac%7B%5Csum_j%20(I_%7Bi%2C%20%5Bj%2C%5Calpha%5D%7D)%7D%7BN_i%7D#0>

*Where*
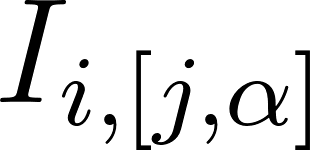
<https://www.codecogs.com/eqnedit.php?latex=I_%7Bi%2C%20%5Bj%2C%5Calpha%5D%7D#0> *is a conditional Indicator Function with*
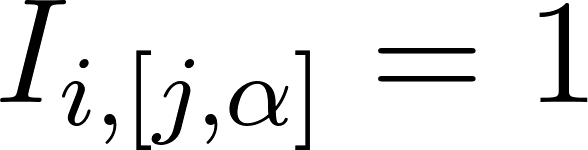
<https://www.codecogs.com/eqnedit.php?latex=I_%7Bi%2C%20%5Bj%2C%5Calpha%5D%7D%3D1#0> *if*
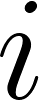
<https://www.codecogs.com/eqnedit.php?latex=i#0> *and*
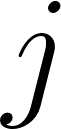
<https://www.codecogs.com/eqnedit.php?latex=j#0> *are adjacent neighbors and cell*
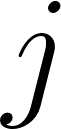
<https://www.codecogs.com/eqnedit.php?latex=j#0> *is in the class*
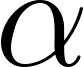
<https://www.codecogs.com/eqnedit.php?latex=%5Calpha#0>*, and*
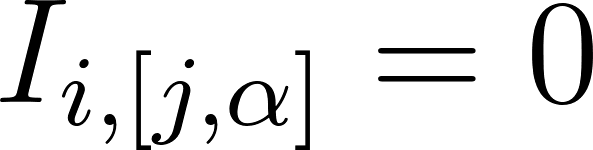
<https://www.codecogs.com/eqnedit.php?latex=I_%7Bi%2C%20%5Bj%2C%5Calpha%5D%7D%3D0#0> *otherwise.*

***Weighted adjacency edge****: if*
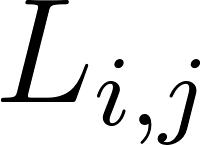
<https://www.codecogs.com/eqnedit.php?latex=L_%7Bi%2Cj%7D#0> *is the weighted indicator function, providing the length of the contact edge between cells*
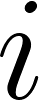
<https://www.codecogs.com/eqnedit.php?latex=i#0> *and*
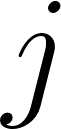
<https://www.codecogs.com/eqnedit.php?latex=j#0>*, the total edge length of contacts for cell*
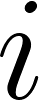
<https://www.codecogs.com/eqnedit.php?latex=i#0> *is*

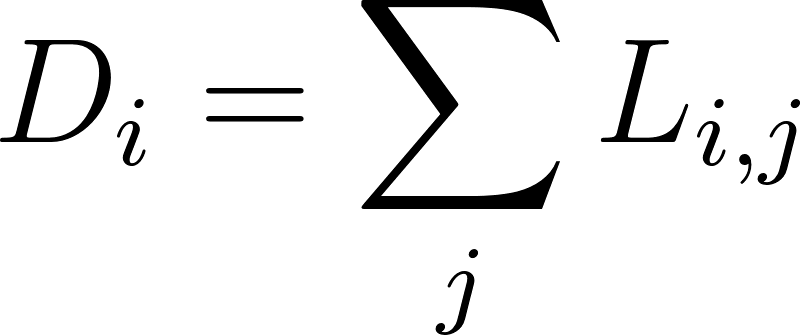
<https://www.codecogs.com/eqnedit.php?latex=D_i%20%3D%20%5Csum_j%20L_%7Bi%2Cj%7D#0>

*The term*
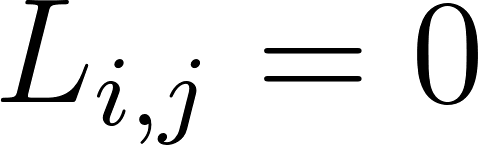
<https://www.codecogs.com/eqnedit.php?latex=L_%7Bi%2Cj%7D%20%3D%200#0> *if*
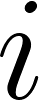
<https://www.codecogs.com/eqnedit.php?latex=i#0> *and*
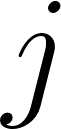
<https://www.codecogs.com/eqnedit.php?latex=j#0> *are not adjacent.*

*Then the fraction of the total edge of a cell that borders with cells of class 𝛂 is given as:*

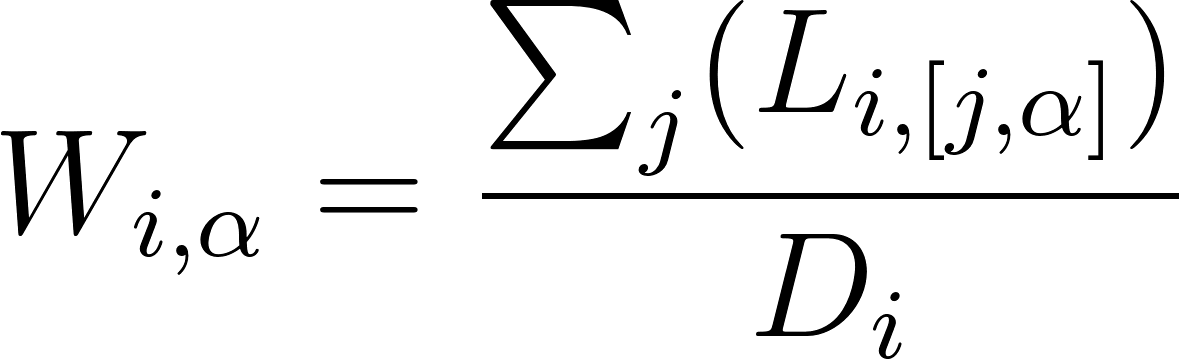
<https://www.codecogs.com/eqnedit.php?latex=W_%7Bi%2C%5Calpha%7D%20%3D%20%20%5Cfrac%7B%5Csum_j%20(L_%7Bi%2C%20%5Bj%2C%5Calpha%5D%7D)%7D%7BD_i%7D#0>

*Where*
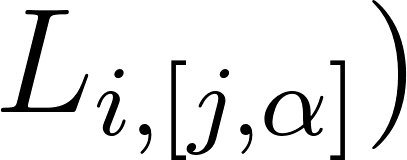
<https://www.codecogs.com/eqnedit.php?latex=L_%7Bi%2C%20%5Bj%2C%5Calpha%5D%7D)#0> *is a conditional weight term defined similarly to the conditional indicator function.*

**Morisita-Horn index:** This second-order metric quantifies the probability that two randomly drawn individuals from two populations
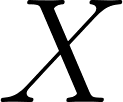
<https://www.codecogs.com/eqnedit.php?latex=X#0> and
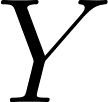
<https://www.codecogs.com/eqnedit.php?latex=Y#0>(typically from different locations in space) are of the same species (1). Therefore, this index estimates whether two populations have similar species composition. The definition for large populations, with
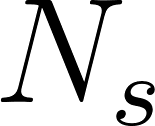
<https://www.codecogs.com/eqnedit.php?latex=N_s#0> the total number of species is:

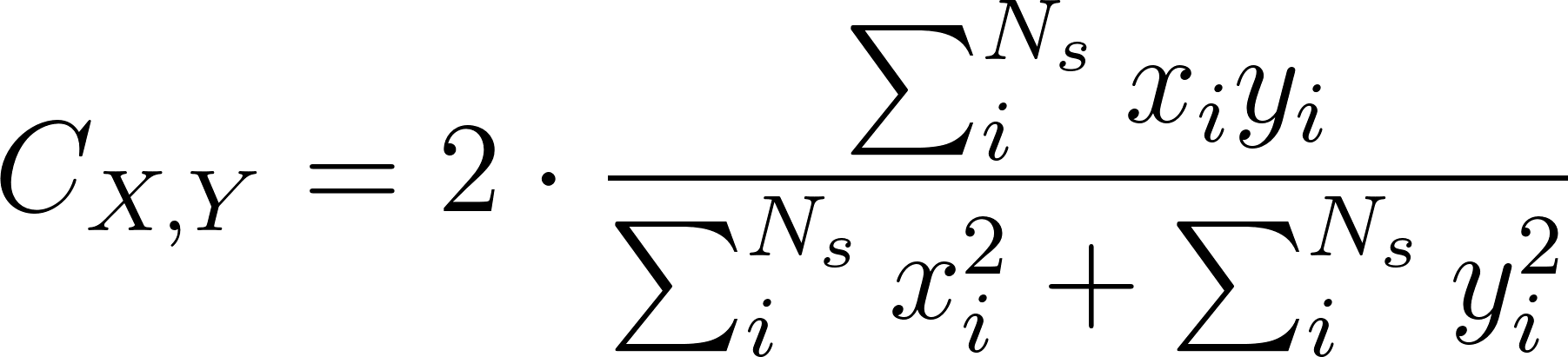
<https://www.codecogs.com/eqnedit.php?latex=C_%7BX%2CY%7D%20%3D%202%5Ccdot%5Cfrac%7B%5Csum_i%5E%7BN_s%7Dx_i%20y_i%7D%7B%5Csum_i%5E%7BN_s%7Dx_i%5E2%20%2B%20%5Csum_i%5E%7BN_s%7Dy_i%5E2%7D#0>

With
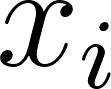
<https://www.codecogs.com/eqnedit.php?latex=x_i#0> and
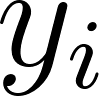
<https://www.codecogs.com/eqnedit.php?latex=y_i#0> the abundances of the
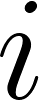
<https://www.codecogs.com/eqnedit.php?latex=i#0>-th species in populations
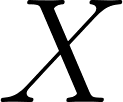
<https://www.codecogs.com/eqnedit.php?latex=X#0> and
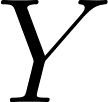
<https://www.codecogs.com/eqnedit.php?latex=Y#0> respectively. This index approaches one if the abundance distributions of
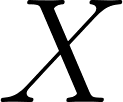
<https://www.codecogs.com/eqnedit.php?latex=X#0> and
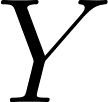
<https://www.codecogs.com/eqnedit.php?latex=Y#0> are similar, and zero if they are mutually exclusive or orthogonal.

**Colocalization index**: This second-order index has a definition similar to the Morisita-Horn index (1). Rather than comparing the species abundance profiles at two locations, this metric quantifies the degree of overlap between the spatial distributions of two classes of points (e.g. species or cell types). This index has been used to study population distribution structures and other applications in ecology (2,3), and more recently for quantifying the spatial colocalization of immune and cancer cells in Breast Cancer (4). The degree of colocalization between two spatial distributions of points:

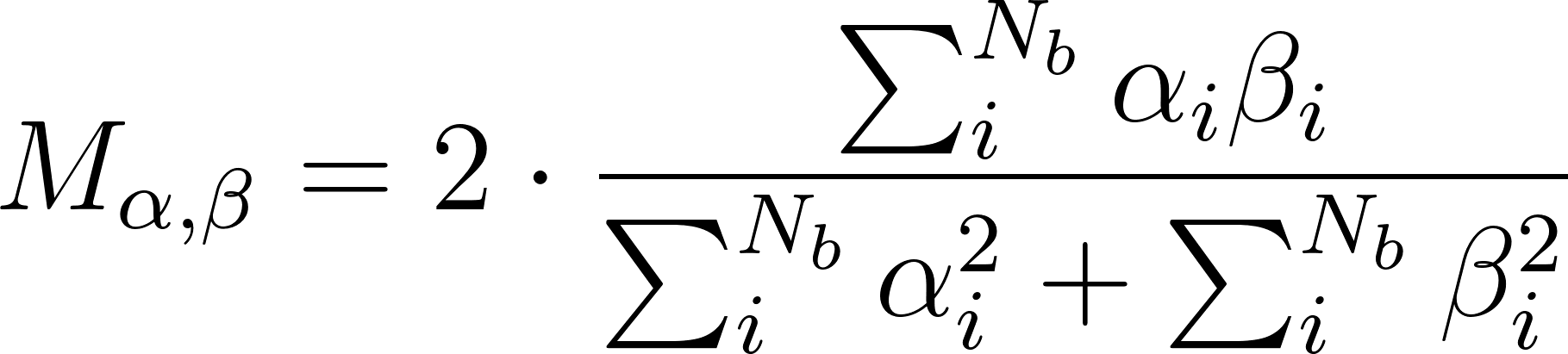
<https://www.codecogs.com/eqnedit.php?latex=M_%7B%5Calpha%2C%5Cbeta%7D%20%3D%202%5Ccdot%5Cfrac%7B%5Csum_i%5E%7BN_b%7D%5Calpha_i%5Cbeta_i%7D%7B%5Csum_i%5E%7BN_b%7D%5Calpha_i%5E2%20%2B%20%5Csum_i%5E%7BN_b%7D%5Cbeta_i%5E2%7D#0>

Where
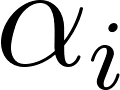
<https://www.codecogs.com/eqnedit.php?latex=%5Calpha_i#0> and
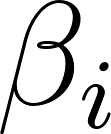
<https://www.codecogs.com/eqnedit.php?latex=%5Cbeta_i#0> are the abundances of points of class
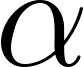
<https://www.codecogs.com/eqnedit.php?latex=%5Calpha#0> and
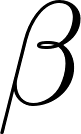
<https://www.codecogs.com/eqnedit.php?latex=%5Cbeta#0> accounted in each location
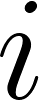
<https://www.codecogs.com/eqnedit.php?latex=i#0> respectively.
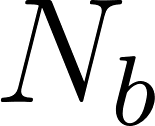
<https://www.codecogs.com/eqnedit.php?latex=N_b#0> is the number of quadrat locations segmenting the space (see diagram).

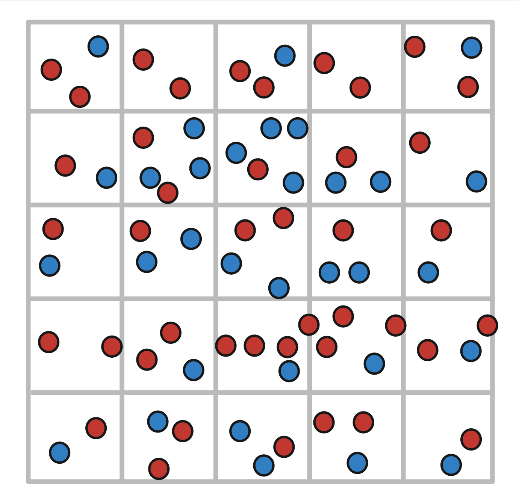

This index is symmetric and approaches a value of one if the two classes have similar spatial distributions and zero if they are spatially segregated from each other.

It is important to note that, as it happens in a
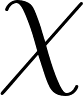
<https://www.codecogs.com/eqnedit.php?latex=%5Cchi#0>-square analysis, this measure is sensitive to low data volume, as it relies on constructing discrete spatial distributions with confidence.

**Nearest Neighbors index**: This second-order index measures the enrichment of nearest neighbors between pairs of point classes. It is based on the standard mean nearest-neighbor distance (MNND) (5–7), implemented in a bivariate form. Let's consider the comparison of the spatial configuration of a particular class of points, called the "test" class, against a reference class "ref" (see diagram).

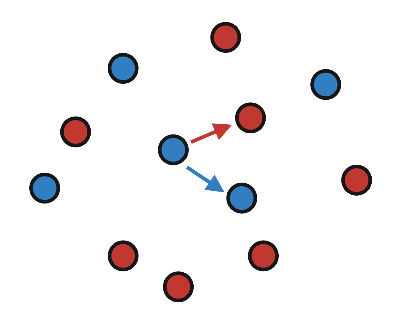

Then if
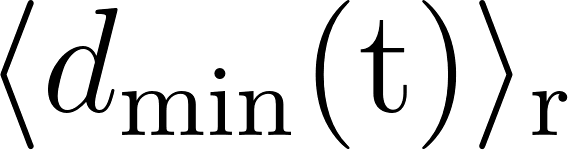
<https://www.codecogs.com/eqnedit.php?latex=%5Clangle%20d_%7B%5Ctext%7Bmin%7D%7D(%5Ctext%7Bt%7D)%5Crangle_%5Ctext%7Br%7D#0> is the MNND of *ref* cells to *ref* cells and
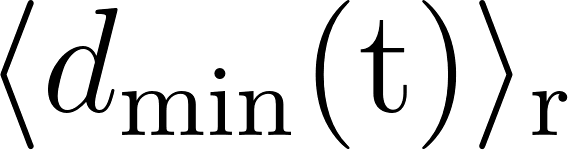
<https://www.codecogs.com/eqnedit.php?latex=%5Clangle%20d_%7B%5Ctext%7Bmin%7D%7D(%5Ctext%7Bt%7D)%5Crangle_%5Ctext%7Br%7D#0> is the MNND of *ref* cells to *test* cells, with
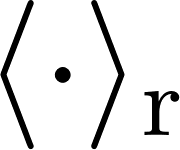
<https://www.codecogs.com/eqnedit.php?latex=%5Clangle%5Ccdot%5Crangle_%5Ctext%7Br%7D#0> the mean overall *ref* cells. The nearest neighbor distance index of *test* cells around *ref* cells is defined as the log ratio:

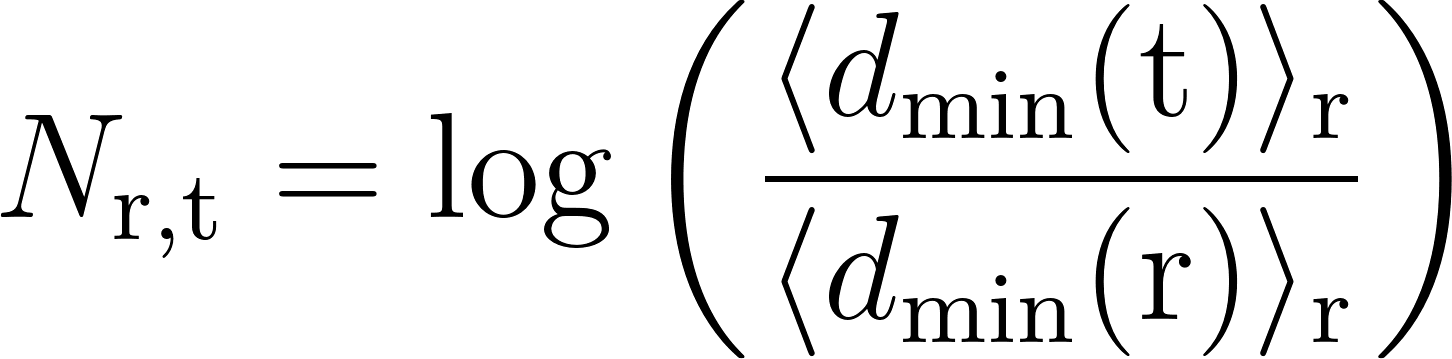
<https://www.codecogs.com/eqnedit.php?latex=N_%7B%5Ctext%7Br%2Ct%7D%7D%20%3D%20%5Clog%5Cleft(%5Cfrac%7B%5Clangle%20d_%7B%5Ctext%7Bmin%7D%7D(%5Ctext%7Bt%7D)%5Crangle_%5Ctext%7Br%7D%7D%7B%5Clangle%20d_%7B%5Ctext%7Bmin%7D%7D(%5Ctext%7Br%7D)%5Crangle_%5Ctext%7Br%7D%7D%5Cright)#0>

This is an asymmetric measure with the following properties:

-
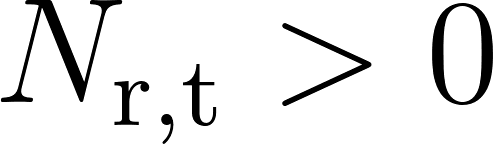
<https://www.codecogs.com/eqnedit.php?latex=N_%7B%5Ctext%7Br%2Ct%7D%7D%20%3E%200#0>: If *ref* and *test* cells are segregated from each other. Thus it is more likely to find a *ref* cell next to a *ref* cell than it is to find a *test* cell.
-
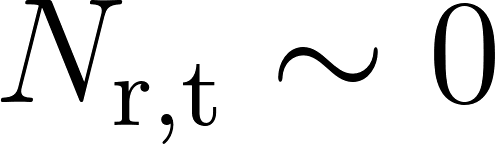
<https://www.codecogs.com/eqnedit.php?latex=N_%7B%5Ctext%7Br%2Ct%7D%7D%20%5Csim%200#0>: If *ref* and *test* cells are well mixed. It is as probable to find a *test* cell next to a *ref* cell as it is to find another *ref* cell.
-
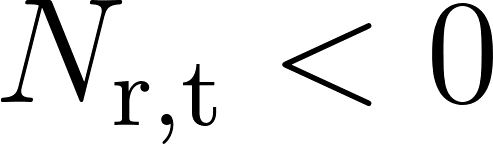
<https://www.codecogs.com/eqnedit.php?latex=N_%7B%5Ctext%7Br%2Ct%7D%7D%20%3C%200#0>: If *ref* cells are individually infiltrated. This means that next to a *ref* cell, it is more likely to find a *test* cell than a *ref* cell. This could happen if ref cells are typically surrounded by *test* cells.

**Ripley’s H index**: This second-order index is derived from the standard Ripley’s
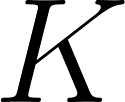
<https://www.codecogs.com/eqnedit.php?latex=K#0> function analysis (8,9). constructed in a bivariate form. It characterizes the relative clumping of spatial distributions of point classes as a function of a description scale. As such, this analysis leads to estimating the system's natural scales and correlation distances (10).

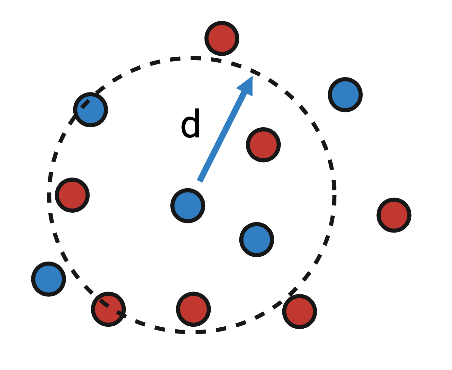

The relative clustering of *test* points around *ref* cells (see diagram) can be estimated by considering the average number of *test* cells in a circle of radius
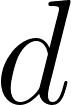
<https://www.codecogs.com/eqnedit.php?latex=d#0> around *ref* cells
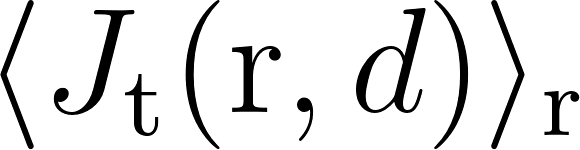
<https://www.codecogs.com/eqnedit.php?latex=%5Clangle%20J_%7B%5Ctext%7Bt%7D%7D(%5Ctext%7Br%7D%2Cd)%5Crangle_%5Ctext%7Br%7D#0>, with the mean taken across all *ref* cells. Then the bivariate Ripley's
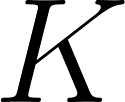
<https://www.codecogs.com/eqnedit.php?latex=K#0> function is this quantity normalized by the density of *test* cells
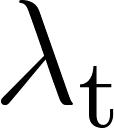
<https://www.codecogs.com/eqnedit.php?latex=%5Clambda_%5Ctext%7Bt%7D#0>:

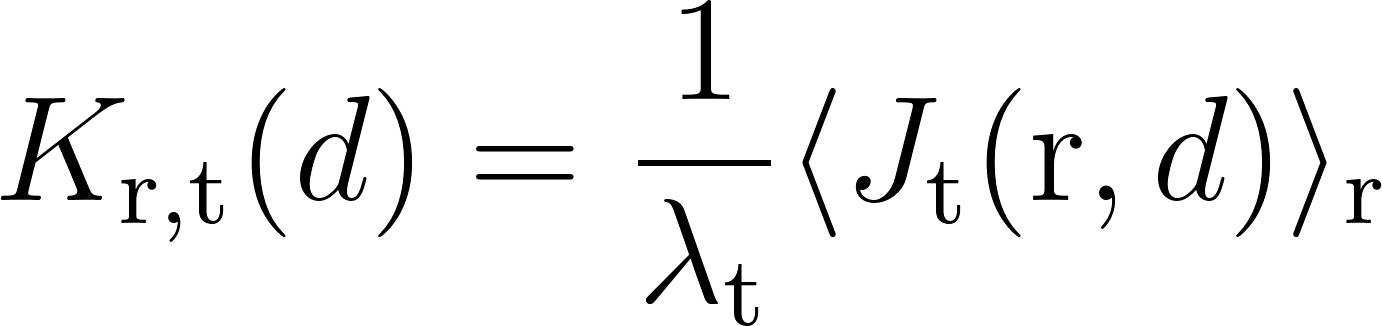
<https://www.codecogs.com/eqnedit.php?latex=K_%7B%5Ctext%7Br%2Ct%7D%7D(d)%3D%20%5Cfrac%7B1%7D%7B%5Clambda_%7B%5Ctext%7Bt%7D%7D%7D%5Clangle%20J_%7B%5Ctext%7Bt%7D%7D(%5Ctext%7Br%7D%2Cd)%5Crangle_%5Ctext%7Br%7D#0>

When the distribution of *test* points is CSR, the expected value is
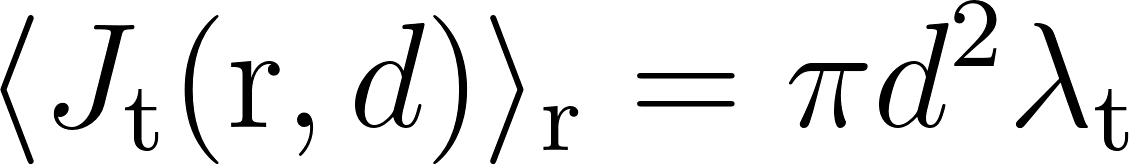
<https://www.codecogs.com/eqnedit.php?latex=%5Clangle%20J_%7B%5Ctext%7Bt%7D%7D(%5Ctext%7Br%7D%2Cd)%5Crangle_%5Ctext%7Br%7D%20%3D%20%20%20%5Cpi%20d%5E2%5Clambda_%7B%5Ctext%7Bt%7D%7D#0>. Then the value of
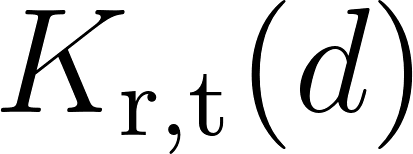
<https://www.codecogs.com/eqnedit.php?latex=K_%7B%5Ctext%7Br%2Ct%7D%7D(d)#0> represents the area of the circle that would contain the observed number of points if they were uniformly distributed (CSR). We define the more convenient function:

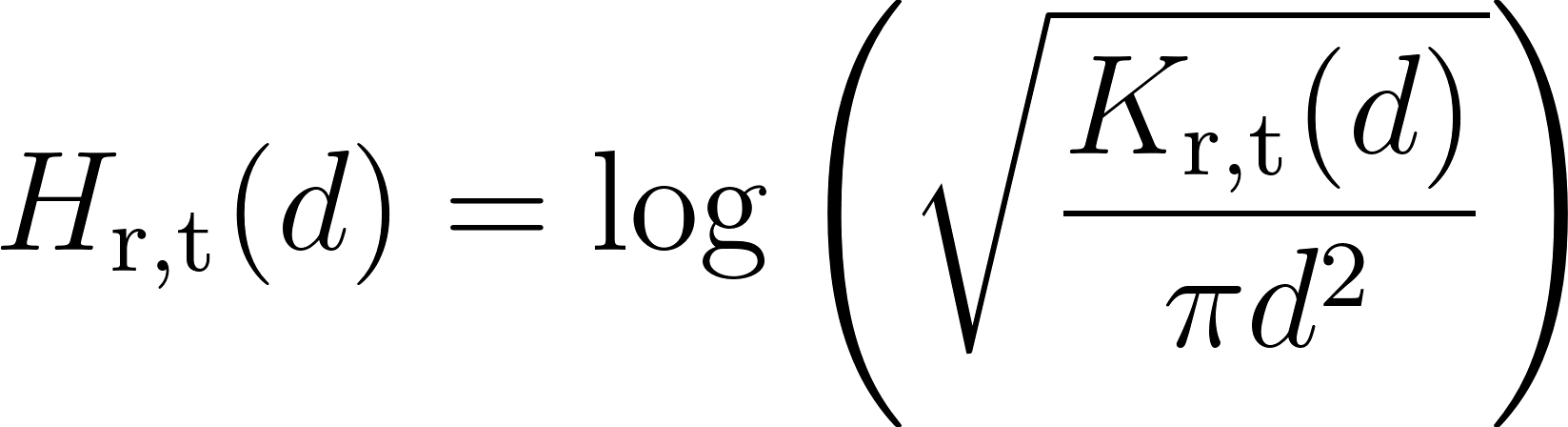
<https://www.codecogs.com/eqnedit.php?latex=H_%7B%5Ctext%7Br%2Ct%7D%7D(d)%20%3D%20%5Clog%7B%5Cleft(%5Csqrt%7B%5Cfrac%7BK_%7B%5Ctext%7Br%2Ct%7D%7D(d)%7D%7B%5Cpi%20d%5E2%7D%7D%5Cright)%7D#0>

This index is not symmetric and has the following properties:

-
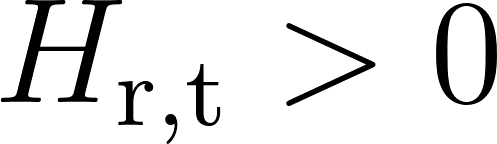
<https://www.codecogs.com/eqnedit.php?latex=H_%7B%5Ctext%7Br%2Ct%7D%7D%20%3E%200#0> if there is a clustering of *test* cells around *ref* cells than expected
-
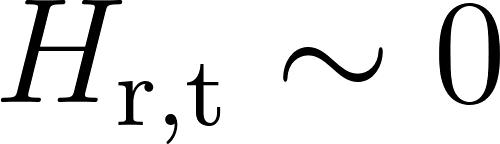
<https://www.codecogs.com/eqnedit.php?latex=H_%7B%5Ctext%7Br%2Ct%7D%7D%20%5Csim%200#0> if there is random mixing between *test* and *ref* cells (CSR)
-
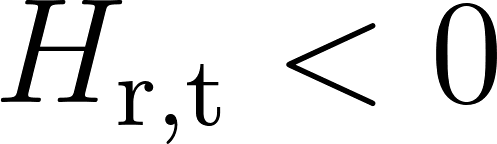
<https://www.codecogs.com/eqnedit.php?latex=H_%7B%5Ctext%7Br%2Ct%7D%7D%20%3C%200#0> if there is random dispersion of *test* cells around *ref* cells

**Getis-Ord Z score**: This is a standard measure of regional enrichment of point density. It consists of a general inferential Z statistic test against the CSR null hypothesis (11–13). Given a point class
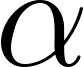
<https://www.codecogs.com/eqnedit.php?latex=%5Calpha#0>, the
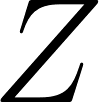
<https://www.codecogs.com/eqnedit.php?latex=Z#0>-score is defined as:

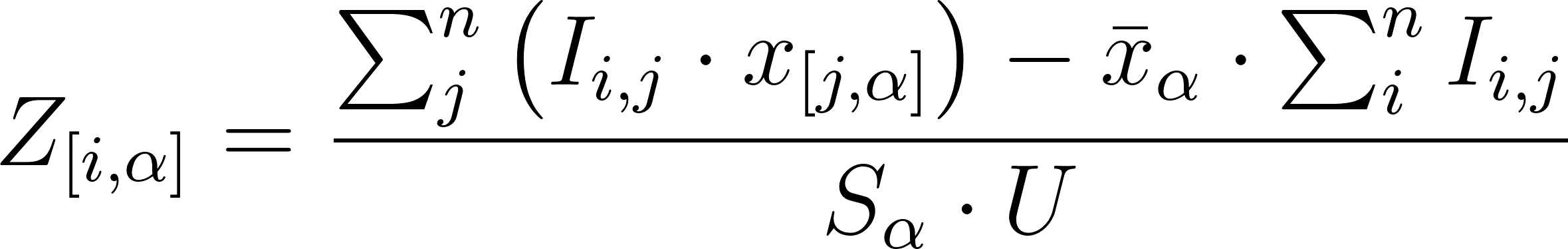
<https://www.codecogs.com/eqnedit.php?latex=Z_%7B%5Bi%2C%20%5Calpha%5D%7D%20%3D%20%5Cfrac%7B%5Csum_j%5En%5Cleft(I_%7Bi%2Cj%7D%5Ccdot%20x_%7B%5Bj%2C%5Calpha%5D%7D%5Cright)%20-%20%5Cbar%7Bx%7D_%5Calpha%5Ccdot%5Csum_i%5En%20I_%7Bi%2Cj%7D%7D%7BS_%5Calpha%5Ccdot%20U%7D#0>

Where
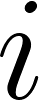
<https://www.codecogs.com/eqnedit.php?latex=i#0> is a discrete location; the term
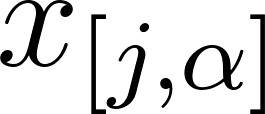
<https://www.codecogs.com/eqnedit.php?latex=x_%7B%5Bj%2C%5Calpha%5D%7D#0> is the abundance of cells of class
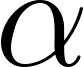
<https://www.codecogs.com/eqnedit.php?latex=%5Calpha#0> in location
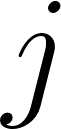
<https://www.codecogs.com/eqnedit.php?latex=j#0>,
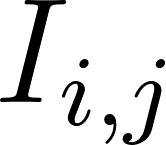
<https://www.codecogs.com/eqnedit.php?latex=I_%7Bi%2Cj%7D#0> is the neighbor indicator function between
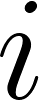
<https://www.codecogs.com/eqnedit.php?latex=i#0> and
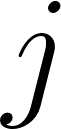
<https://www.codecogs.com/eqnedit.php?latex=j#0> (defined as 1 if
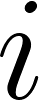
<https://www.codecogs.com/eqnedit.php?latex=i#0> and
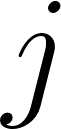
<https://www.codecogs.com/eqnedit.php?latex=j#0> are neighbors and 0 otherwise),
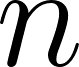
<https://www.codecogs.com/eqnedit.php?latex=n#0> is the total number of locations,
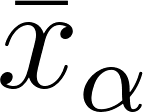
<https://www.codecogs.com/eqnedit.php?latex=%5Cbar%7Bx%7D_%5Calpha#0> is the mean abundance of cells of class
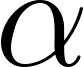
<https://www.codecogs.com/eqnedit.php?latex=%5Calpha#0> and the dispersion terms:

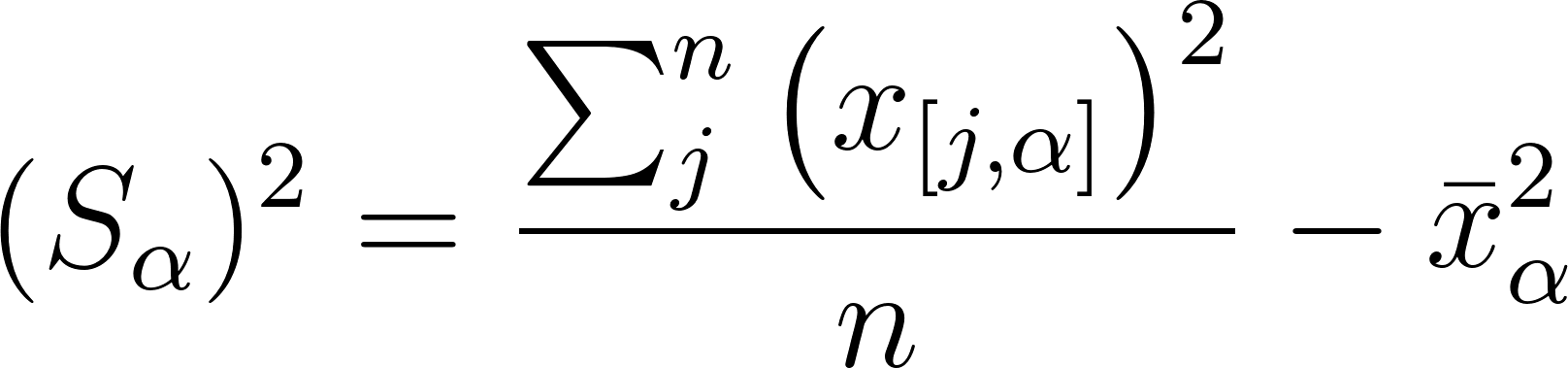
<https://www.codecogs.com/eqnedit.php?latex=%20(S_%5Calpha)%5E2%20%3D%20%7B%5Cfrac%7B%5Csum_j%5En%20%5Cleft(x_%7B%5Bj%2C%5Calpha%5D%7D%5Cright)%5E2%7D%7Bn%7D%20-%20%5Cbar%7Bx%7D_%5Calpha%5E2%7D#0>

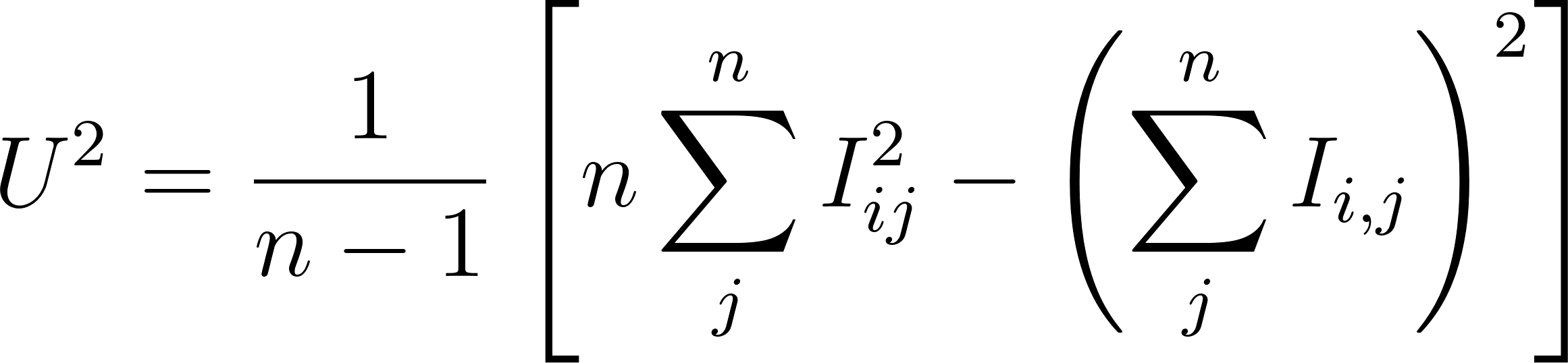
<https://www.codecogs.com/eqnedit.php?latex=%20U%5E2%20%3D%20%7B%5Cfrac%7B1%7D%7Bn-1%7D%5Cleft%5Bn%5Csum_j%5En%20I_%7Bij%7D%5E2%20-%20%5Cleft(%5Csum_j%5EnI_%7Bi%2Cj%7D%5Cright)%5E2%5Cright%5D%7D#0>

A corresponding p-value can be calculated from
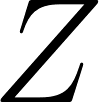
<https://www.codecogs.com/eqnedit.php?latex=Z#0>. If  is positive and statistically significant, the observed index value in the location  is larger than expected from a CSR assumption, thus indicating a hot spot of point abundance. In the same way, if the score is significant and negative it indicates a depletion of points in that location.

**Localized point statistics:** The two-dimensional convolution smoothing function is defined as (14):

where  is the kernel weight,  is the spatial function to be smoothed, and we define the term as the convolution operator. With this:

- *Colocalization index* between classes with abundances  and  for  and  cells correspondingly, is measured in each region as:

- *Nearest Neighbor Index* of *test* cells around *ref* cells is calculated locally as:

With  the local mean of the function  over *ref* cells in the local region, and  is the number of *ref* cells in the local region.

- *Ripley’s H index* of *test* cells around *ref* cells is calculated locally as:

Where  is Ripley’s radius, and  is the local density of *test* cells. The local mean is calculated over all ref cells in the neighborhood as:

.

In field ecology, quadrats are typically used to quantify small regions and produce spatial profiles across a landscape. Then, each neighborhood is binned down into even smaller regions to construct a spatial distribution profile of cell abundances. Using this same principle, our methodology uses a bandwidth parameter to set a scale length to coarse-grain the landscape mosaic in terms of local properties of cell abundance and uniformity in their spatial distribution (which we call “mixing”). In each quadrat partitioning the landscape around location, we calculate the abundance  of each cell type , as well as the value of a mixing index  for each cell type, defined as:

Which is calculated over  “sub-quadrats” in the neighborhood of ,  are the abundances in each of those sub-quadrats, and

This is a univariate form of the Morisita-Horn index comparing the observed spatial profile with an array of the same size and a uniform CSR distribution = constant. This score is a simple way to account for the degree of mixing (uniformity) of cells in a sample. A value  means that the sample is highly segregated (ie. the variance across sub-quadrats is large) and a value  means that all sub-quadrats have very similar count values and thus, cells are uniformly distributed across the quadrat.

LME classes are defined in three general categories for each cell type: (B) Bare environments are those with very few cells, regardless of mixing. (S) Segmented environments are those in which cells are clustered together; they have moderate to high abundance and low mixing. (M) Mixed environments are those where cells are mixed uniformly; they have moderate to high abundance and high mixing. Because the mixing score is sensitive to low abundance, medium abundance levels are considered "mixed" (M) if the mixing is medium, as mixing is biased to lower values with lower abundances. With is scheme, (B, S, M) codes are assigned for each individual class, forming categories like: `BBB, BBS, BBM …. MMM`. Each of these categories represents a basic type of Local Micro-Environment that is consistent across the entire study and can be used to define ecological phenotypes in the TLA.

Fragmentation statistics are summarized at different hierarchical levels:

**Patch-level metrics**: These are divided into six different statistics, calculated over individual patches for each class type:

**Area**: The total area spanned by -th patch of class  (i.e. patch ) is:

**Perimeter**: The smoothed boundary length patch  is:

**Edge**: The length of the edge boundary between two neighboring patches  and  is:

**Perimeter-area ratio**: The ratio of perimeter and area for a patch. This is a measure of the morphological complexity of patch :

Note: this metric is scale-dependent as the perimeter grows linearly with the length scale and the area grows as the square.

**Shape index**: Measures patch  shape complexity as the ratio of the side of a square with the same perimeter and to the side of a square with the same area as the patch. This metric is scale-corrected:

- : The patch is close to a simple square shape, or minimal perimeter for a given area.
- : As the shape becomes more irregular, the shape index increases. For irregular shapes, the perimeter increases more than the area does, leading to larger values for the shape index.

**Fractal Dimension**: Measures the patch shape complexity as an approximation to the fractal dimension:

- : The patch is close to a simple square shape, or minimal perimeter for a given area.
- : The patch boundary is a complex, plane-filling, shape. Hence the perimeter has a fractal dimension >1.

**Euclidean nearest neighbor**: The distance to the nearest neighboring patch of the same class using the Euclidean distance:

Where  is the shortest edge-to-edge distance from patch  to . This measure provides a quantification of the typical proximity between patches of the same class.

**Class-level and landscape-level metrics**: These metrics are aggregated by summarizing patch-level statistics over all the patches in the same class and all the classes across the landscape, accordingly:

**Number of patches**: The number of patches of class  is  and the total number of patches is:

**Area**: Total area in pixels covered by a class  and the total area are:

**Proportion of Landscape**: The proportion of the landscape area covered by a certain class :

This measure quantifies the importance or prevalence of each class of habitat in the landscape.

**Patch Density**: The relative density of patches per unit area for class , and the total patch density:

This measure is a quantification of both the prevalence of a class in the landscape, but it is also an indicator of fragmentation: between two landscapes with similar values of , the one with a larger  value is more fragmented than the other. This is a more general index than the patch area.

**Patch index**: the proportion of the landscape comprised by the largest patch in class :

Similarly, the total patch index is the proportion of the landscape comprised by the largest patch:

**Total edge**: the edge length of class  and total edge in the landscape:

**Edge Density**: The edge density for class  and the total edge density are normalized by the total landscape area:

This metric is a normalized measure of the amount of contact between patches, which is related to both the level of complexity of the patches' morphologies and their level of fragmentation.

**Landscape shape index**: This index is a metric of class aggregation that provides a normalized measure of edginess adjusted for the size of the landscape:

This is another measure of fragmentation that accounts for the level of complexity of patch boundaries in the landscape.

**Contagion:** also called Interspersion, is a landscape-level measure that accounts for the probability of adjacent locations belonging to the same class. It quantifies how aggregated or dispersed different classes are according to the landscape's conformation.

Where  is a normalized indicator function term that accounts for patch adjacencies, and  is the total number of classes.

- means that the landscape is highly disaggregated (cells are spread out, and each cell belongs to a different class), which means that it is more likely that the adjacent cells belong to different classes.
- indicates that the adjacent cells are more likely to belong to the same class. In this scenario, the landscape is more aggregated.

**Shannon Diversity Index**: This metric quantifies the diversity of a landscape based on the number and abundance of different classes.

With  the normalized frequency of class . The value of  increases as the entropy increases (number of classes increases) and it is close to 0 if the landscape is homogenous.
